# Representation of sex-specific social memory in ventral CA1 neurons

**DOI:** 10.1101/2024.04.01.587523

**Authors:** Akiyuki Watarai, Kentaro Tao, Teruhiro Okuyama

## Abstract

Recognizing familiar individuals is crucial for adaptive social interactions among animals. However, the multidimensional nature of social memory encompassing sexual information remains unelucidated. We found that neurons in the ventral CA1 region (vCA1) of the mouse hippocampus encoded the identities and social properties, specifically sex and strain, of familiar conspecifics by using both rate and theta-based temporal coding. Optogenetic reactivation of social memories of females, but not males, induced place preference. Ablation of the upstream hippocampal dorsal CA2 region (dCA2) or the medial amygdala (MeA) disrupted the representation of sex and the sexual dichotomy of social memory valence. Selective reactivation of overlapping neural populations of distinct female social memories representing the female sex was sufficient to induce preference. Thus, vCA1 neurons employ dual coding schemes to represent the identities and social properties of familiar conspecifics as a cohesive memory.

**One-Sentence Summary:** Social memory in the mouse ventral hippocampus maps the identities and social properties of familiar conspecifics.

## Main Text

The hippocampus plays a crucial role in the integration of multimodal information into a conceptualized cognitive map, which is fundamental for recalling specific experiences and guiding behaviors (*1*). Social memory is a key component of episodic and declarative memory, that enables the recognition of familiar individuals. Extensive animal studies, particularly those using rodent models, have underscored the crucial role of the hippocampus in processing social memory. Located along the hippocampal septotemporal axis, the dorsal CA2 region (dCA2) specializes in encoding social stimuli (*2*, *3*), whereas the ventral CA1 region (vCA1), which receives projections from dCA2 (*4*), is responsible for storing memories of familiar conspecifics (*5*) through the temporal coordination of neuronal population activities (*6*). Notably, a recent study in rats revealed that neuronal responses in the vCA1 to familiar conspecifics showed greater variability among individual females than the responses among males (*7*), suggesting a potential sexually dichotomous component in hippocampal social memory. Despite significant advances, the neural mechanisms by which hippocampal networks assemble various elements of social information, such as genetic and physical traits (including sex), into cohesive social memories remain largely unelucidated.

### Nonlinear mixed representation of familiar conspecifics by vCA1 neurons

To investigate how vCA1 neurons respond to different familiar conspecifics, we developed a behavioral paradigm in which mice (*n* = 6 adult C57BL/6 male mice) were implanted with a high-density silicon probe in the vCA1 (Fig. 1A), explored the arena, and interacted with four different familiarized conspecifics that comprised either a female or a male mouse of the BALB/c or C3H strains (BF, BM, CF, and CM; Fig. 1B and fig. S1, A and B). An analysis of the interaction duration (fig. S1, C to E) and head direction (fig. S2) confirmed that the subjects were engaged with the social stimulus, particularly during nose-poking into the social chamber, when neuronal spikes were frequently observed (fig. S3). Examination of the vCA1 neural activity revealed the presence of putative pyramidal cells (fig. S4A) activated by the interaction with a specific mouse stimulus (Fig. 1, C and D; additional examples are presented in fig. S4B) and its associated properties, including sex and strain (fig. S4C).

**Fig. 1.**
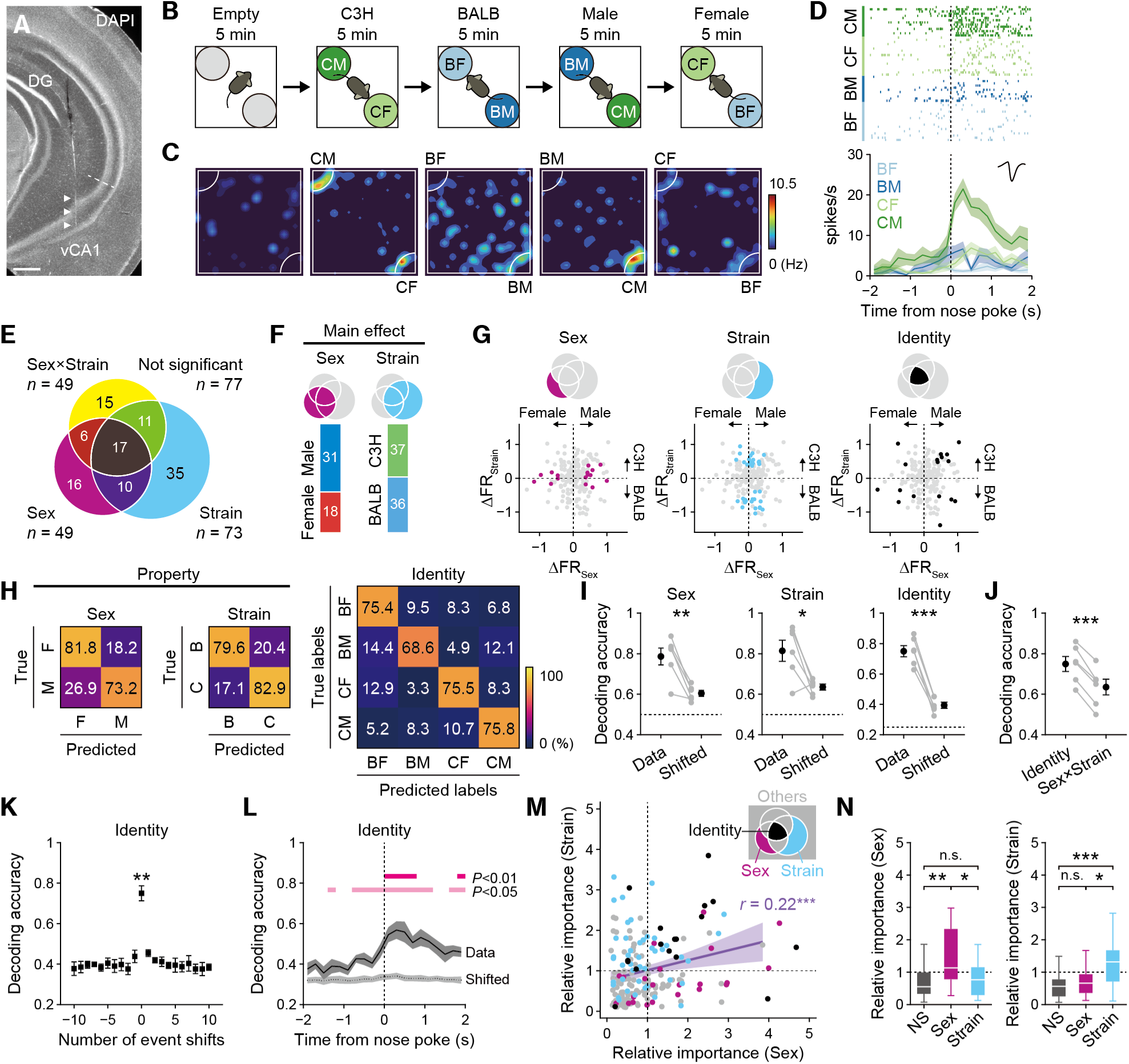
Ventral CA1 neurons encode the identity and properties of social memory. (**A**) Recording site. Scale bar, 200 µm. (**B**) Paradigm for the four-mouse social interaction test. BF, BM, CF, and CM denote BALB/c females, BALB/c males, C3H females, and C3H male mice, respectively. (**C**) Firing rate heatmaps of a representative CM neuron. Spikes could be assigned to the area within the social chambers since we tracked the tip of the nose as the reference body point of the subject. For more detail, see fig. S3. (**D**) Peri-event time histogram and raster plots aligned to the nose-poke onset for the same neuron as in (C). (**E**) The Venn diagram illustrates the results of two-way ANOVA for all recorded putative pyramidal cells (*n* = 187). Significance was determined at *P* < 0.05. (**F**) Target preferences of cells with a significant main effect of sex or strain. (**G**) Distribution of sex (*n* = 16), strain (*n* = 35), and identity (*n* = 17) cells on a 2-D map showing differences in z-scored firing rates by sex and strain of social stimuli. (**H**) Confusion matrices for decoding sex (left), strain (center), and social identity (right), averaged over the subjects (*n* = 6 mice). The matrices were normalized by row to represent the true positive and false negative rates. (**I**) Decoding accuracy of classifiers constructed using real or shifted event labels. \**P* < 0.05, \*\**P* < 0.01, \*\*\**P* < 0.001, paired *t* test. (**J**) Comparison of the decoding accuracy of identity classifiers with the linear combination of decoding accuracy of sex and strain classifiers. \*\*\**P* < 0.001, paired *t* test. (**K**) Decoding the accuracy of the classifier with real data (shifts = 0) and event-shifted labels (shifts = ±10 events) for identity. \*\**P* < 0.01 for shifts = 0 vs. ±1, Tukey–Kramer multiple comparisons. (**L**) Decoding accuracy of identity over time (mean ± SEM, *n* = 6 mice). Significance was assessed by Tukey–Kramer multiple comparisons. (**M**) Scatter plot of the relative importance of sex decoding versus strain decoding and their correlation (*r* = 0.22, *P* < 0.002, permutation test) for all recorded pyramidal cells (*n* = 187). Red dots represent sex cells (*n* = 16), blue dots represent strain cells (*n* = 35), black dots represent social identity cells (*n* = 17), and gray dots indicate all other pyramidal cells (*n* = 136). (**N**) Comparison of relative importance between sex and strain cells. NS, non-social. \**P* < 0.05, \*\**P* < 0.01, \*\*\**P* < 0.001, Wilcoxon rank-sum test.

To classify neurons according to their response selectivity to social stimuli, we performed a two-way ANOVA (*8*, *9*) on firing rates during social interactions, with sex and strain as main factors and their interaction term. This generated eight subpopulations with distinct activity patterns in response to social stimuli (Fig. 1E). Among all recorded putative pyramidal cells, 26% (49/187) and 39% (73/187) showed significant main effects of sex and strain, respectively. In contrast, 26% (49/187) encoded a significant nonlinear interaction between sex and strain. Both groups of cells with a significant main effect of sex or strain comprised a statistically comparable number of neurons preferring either females or males and either BALB/c or C3H strains (Fig. 1F). Cells with only a significant main effect of either sex or strain were defined as social property cells, while cells with significant main effects of both factors and a significant interaction term were defined as social identity cells (Fig. 1G and fig. S4, D and E). The overall firing rates were comparable between social property cells and social identity cells (fig. S4F). The probe placement successfully targeted vCA1 while sparing dCA2 (fig. S5, A and B), resulting in a uniform distribution of social cells across the anteroposterior axis of the ventral hippocampus (fig. S5C). By contrast, targeting dorsal CA1 (dCA1) yielded significantly fewer neurons responsive to sex, strain, and identity (*n* = 3 mice across 4 sessions; fig. S6).

### Multiplexed coding of social identity and properties

Based on the neuronal activity during social interactions, we trained a set of linear classifiers (Fig. 1H). All classifiers achieved significantly higher decoding accuracies than the null models, which were constructed by circularly shifting event labels (fig. S7A), at both the group level (fig. 1I) and individual subject level (fig. S7B). The null models constructed by random shuffling yielded the same results (fig. S7, E and F). The linear combination of classifiers for sex and strain did not incorporate a high decoding accuracy of the identity classifier (Fig. 1J), suggesting that nonlinear social memory information is embedded in vCA1 population activity. Although the social interaction sequences exhibited a nonrandom pattern characterized by consecutive staying and switching events, the classifier constructed using real labels significantly outperformed the surrogate classifier, constructed using labels shifted by only one (Fig. 1K and fig. S7C). Corresponding to the transient rise in firing rate during social interactions (Fig. 1D), we observed that the decoding accuracy of the identity classifiers increased over time as the subject mice approached the stimulus and poked their noses into the social chamber (Fig. 1L).

This was the case for both the sex and strain classifiers (fig. S7D). To ascertain whether specific subpopulations of neurons encode social properties, we analyzed the weights assigned to each neuron using classifiers trained to decode both sex and strain (Fig. 1M and fig. S8A). A significant positive correlation between sex and strain was observed across all recorded pyramidal cells (Fig. 1M), which was also evident in the simultaneously recorded pyramidal cells from four of the six subjects (fig. S8B). These findings suggest that a subset of vCA1 neurons modulate the encoding of mixed social memory information. A comparison of the relative importance based on single-cell classification revealed that sex cells were more crucial for decoding sex than strain cells and non-social cells, which were equally below the average importance. Strain cells were more influential for decoding strain than sex cells and non-social cells, both of which showed comparable below average importance (Fig. 1N).

To assess whether the representation of social properties conveyed generalized information across conditions, we trained a linear classifier using pseudo-population data derived from a subset of conditions (e.g., interactions with female mice only) to discriminate a pair of social stimuli (BF and CF). Then, we tested its decoding accuracy using data from the complementary conditions (BM and CM). Both the sex classifiers across the strains (fig. S9, A to D), and strain classifiers across the sexes (fig. S9, E to H) exhibited a significantly higher cross-condition generalization performance (CCGP) (*10*) than their respective null models (fig. S9, B and F).

The strain classifier trained on males was less accurate than that trained on females (fig. S9G), which is in line with a previous study that demonstrated that vCA1 neuronal responses to females exhibited greater diversity among individual females than among males (*7*).

### Theta-based temporal coding of social memory information

A detailed analysis of the neuronal firing patterns showed that a social identity cell exhibited phase locking of spike timings with the hippocampal theta trough (fig. S10A), which corresponded to a transient increase in the theta-band spiking activity that was coherent with theta oscillations (fig. S10B). A comparison of spike-LFP coherence at the onset of social interactions showed that social cells exhibited greater phase locking to the theta rhythm than non-social cells (fig. S10, C to E). Social and non-social cells displayed distinct theta phase preferences, with the former favoring the trough and the latter spanning from the ascending phase to near the peak (Fig. 2, A to C). Cells encoding social identity and properties showed a preference for theta troughs. However, social identity cells exhibited a more pronounced concentration around the trough (fig. S10F). During interactions with preferred versus non-preferred social stimuli for each cell, only social property cells demonstrated a shift toward earlier theta phases, which was indicative of differential phase locking (Fig. 2, D and E). Although interactions with preferred stimuli led both cell types to favor the theta trough, in contrast to social identity cells, social property cells uniquely preferred later theta phases when faced with nonpreferred stimuli (Fig. 2F).

**Fig. 2.**
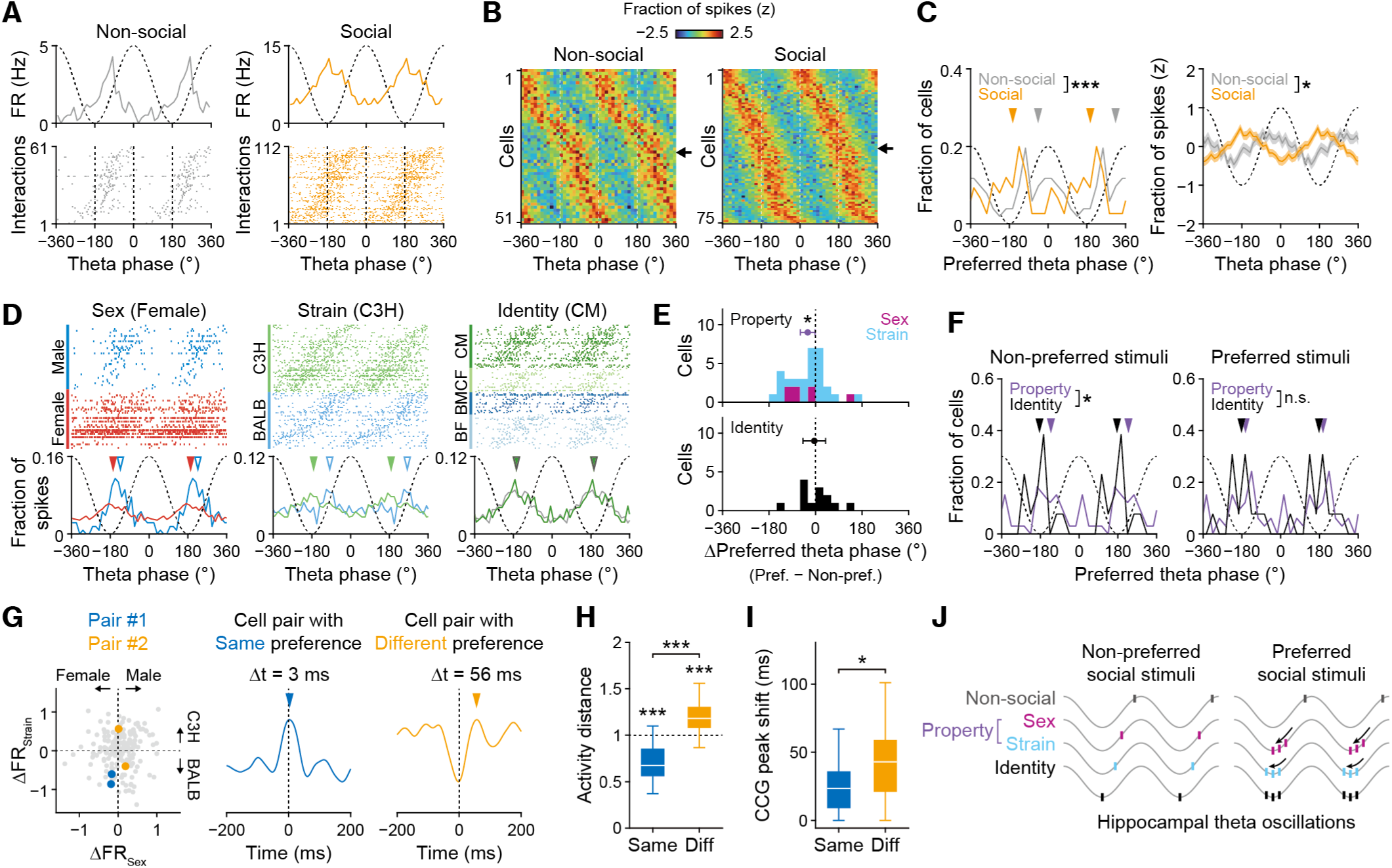
Theta phase-specific encoding of social identity and its properties. (**A**) Raster plots and spike theta phase distributions for an example non-social cell and an example social cell. (**B**) Spike theta phase distributions for non-social cells (*n* = 51) and social cells (*n* = 75), organized by preferred phases. Only cells with significant theta modulation (*P* < 0.01, Rayleigh test) were considered. Arrows indicate the example cells shown in (A). (**C**) Left, distribution of the preferred theta phases for social and non-social cells. The arrowheads in (C) and (F) indicate the average preferred phase for each group. \*\*\**P* < 0.001, Watson–Williams test. Right, mean z-scored proportions of spike theta phases for social and non-social cells. \**P* < 0.05, permutation test. (**D**) Raster plots and spike theta phase distributions from left to right for an example sex cell (female preferred), an example strain cell (C3H preferred), and an example identity cell (CM preferred). Spike raster rows for each social interaction were organized according to the preferred theta phases of the trial. The mean theta phases of spikes for the preferred and non-preferred social stimuli are represented by filled and empty arrowheads, respectively. (**E**) Distribution of preferred spike phase differences when interacting with preferred versus nonpreferred social stimuli. \**P* < 0.05, one-sample test for the mean angle. Error bars represent a 95% confidence interval. (**F**) Distribution of preferred theta phases for social property cells and social identity cells when interacting with preferred or nonpreferred social stimuli. \**P* < 0.05, Watson–Williams test. (**G**) Representation of two cell pairs on a 2-D map showing differences in z-scored firing rates by sex and strain of social stimuli. (**H**) Comparison of pairwise distances in social neuronal activity between cell pairs with the same (*n* = 44) and different (*n* = 21) social property preferences. \*\*\**P* < 0.001, Wilcoxon rank-sum test. (**I**) Comparison of CCG peak shifts between cell pairs with the same versus different social property preferences. \**P* < 0.05, Wilcoxon rank-sum test. (**J**) Schematic summary of the rate and temporal coding strategy in vCA1 to represent social identity and properties.

To further elucidate the temporal coding structure of vCA1 activity during social interactions, we compared the pairwise distances of neuronal activity and time lags between the spikes of the two corresponding cells at theta timescales (fig. S11). Only cell pairs exhibiting significant theta modulation in spike cross-correlograms (CCGs) were included (*P* < 0.01, permutation test; fig. S11A), with a significantly higher proportion of social cells (365/1304 pairs, 28.0%) than that of non-social cells (33/517 pairs, 6.0%) (fig. S11B). Among pairs of social cells, but not non-social cells, a significant positive correlation was observed between the activity distance and temporal peak shift of the smoothed CCGs, which was more pronounced when only pairs of either two sex cells or two strain cells were analyzed (fig. S11D). Furthermore, we classified pairs of social property cells based on whether the two cells had the same preference for either sex or strain of social stimulus (Fig. 2G). As expected, the activity distances between cell pairs with identical social property preferences were significantly shorter than those between pairs with different social property preferences (Fig. 2H). Notably, the temporal shifts in the CCG peaks were shorter for pairs with the same preference than for those with different preferences (Fig. 2I). These results underscore a rate and temporal coding strategy in neurons that represent social identities and properties, with a notable divergence of theta phase precision in social property cells (Fig. 2J).

### Reactivation of female social memory induces place preference

To capture the representation of sex in social memory, we utilized recently developed techniques for observing and manipulating engrams (*11*, *12*). Specifically, we combined activity-dependent cell labeling using a virus-mediated system with immunohistochemical cell labeling for immediate early gene expression. Mice were injected with two adeno-associated viruses (AAVs) into the vCA1: one expressing the tetracycline transcriptional activator (tTA) driven by the *c-fos* promoter (AAV2-c-fos:tTA) and the other expressing histone-GFP regulated by the tetracycline response element (TRE) when doxycycline (Dox) was absent from the diet (AAV5-TRE:HTG) (Fig. 3A). This enabled the genetic labeling of neurons that were activated during the first social interaction. For the second social interaction, we used c-Fos immunostaining to visualize activated neurons (Fig. 3B). The mice were divided into three groups as follows: mice that interacted first with female BALB/c and then with female C3H (FF), mice that interacted with female BALB/c and then with male C3H (FM), and mice that interacted with male BALB/c and then with male C3H (MM) (Fig. 3C). Although all groups displayed a comparable number of active neurons during both the first (HTG^+^) and second (c-Fos^+^) interactions, the group that interacted with social stimuli of different sexes (FM) showed a significantly lower number of neurons that were double-labeled with HTG and c-Fos as well as a reduced proportion of double-positive neurons among the HTG^+^ neurons (Fig. 3, D and E). These results suggest that distinct social memory neurons encode the representation of sex which corroborates the results of the neural recordings (Fig. 1).

**Fig. 3.**
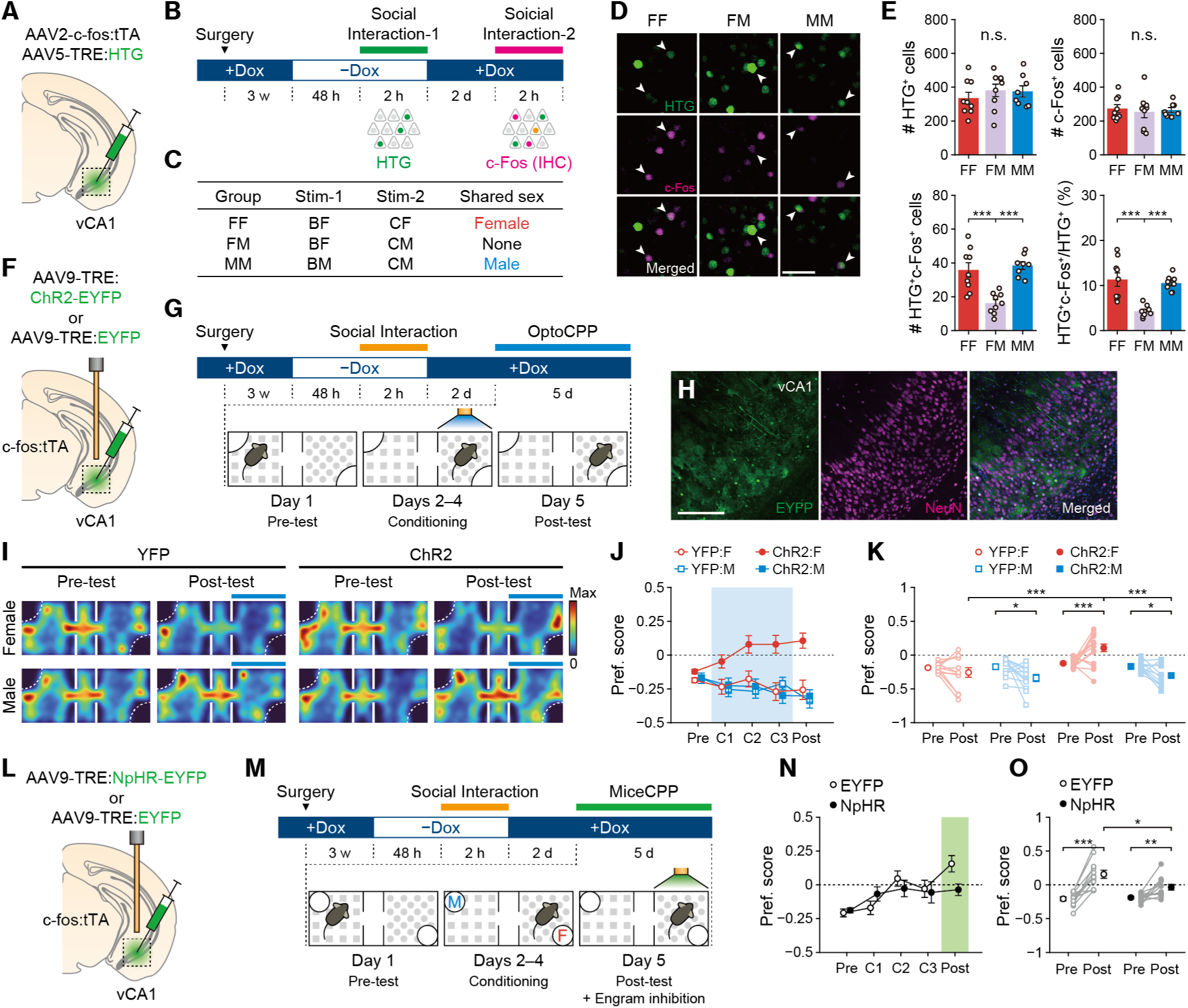
Reactivation of female social memory in vCA1 exerts a positive valence. (**A**) Schematic representation of virus-mediated activity-dependent cell labeling targeting vCA1. (**B**) Experimental paradigm for activity-dependent double cell labeling. Neurons activated by the first social interaction were labeled by HTG expression, whereas those activated by the second interaction were detected by c-Fos expression. (**C**) Labeling paradigm for the three behavioral groups. BF, BM, CF, and CM represent BALB/c females, BALB/c males, C3H females, and C3H males, respectively. (**D**) Representative images of labeled vCA1 neurons. Scale bar, 50 µm. (**E**) Number of HTG^+^ cells, number of c-Fos^+^ cells, number of HTG^+^c-Fos^+^ double-positive cells, and ratio of double-positive cells to HTG^+^ cells. (**F**, **G**) Schematic (F) and experimental paradigm (G) for the optogenetic conditioned place preference (optoCPP) test targeting vCA1. (**H**) Representative images of vCA1 showing the EYFP expression. Scale bar, 100 µm. (**I**) Occupancy maps averaged across subjects before (Pre-test) and after (Post-test) optogenetic conditioning. Filled blue bars indicate the stimulated side during conditioning (i.e., the nonpreferred side during the Pre-test). (**J**) Temporal dynamics of preference scores during conditioning (*n* = 12–16 mice per group). (**K**) Comparison of preference scores between the Pre-test and Post-test. (**L**) Schematic representation of virus-mediated labeling of the social memory engram with an inhibitory opsin. (**M**) Experimental paradigm for conditioned place preference test with live mice as stimuli. (**N**) Temporal dynamics of preference scores during conditioning (*n* = 12 mice per group). F: female, M: male. (**O**) Comparison of preference scores between the Pre-test and Post-test. \**P* < 0.05, \*\**P* < 0.01, \*\*\**P* < 0.001; n.s., not significant. Error bars represent SEM.

To examine whether the social memory of a conspecific carried a specific valence, we employed an optogenetic conditioned place preference (optoCPP) test (*13*) by injecting AAV9-TRE:ChR2-EYFP or AAV9-TRE:EYFP into the vCA1 of c-fos:tTA mice and implanted optic fibers directly above the injection site (Fig. 3F). When the subject mouse was off-Dox for 48 h (Fig. 3G), a 2-hour exposure to either a female or male C3H mouse resulted in sparse transgene expression (Fig. 3H) with comparable cell counts (fig. S12A). Light stimulation of the social memory engram linked to a female conspecific, but not a male, induced a positive reinforcing effect on the post-test (day 5), as well as a real-time place preference during conditioning (days 2−4; Fig. 3, I to K). This sexually dichotomous effect on social memory reactivation was observed when female and male BALB/c mice were used to label the engram (fig. S12, B and C), suggesting that the representation of the female sex had a positive valence regardless of the strain of the social stimulus. When female mice were used as subjects, the activation of female social memory engram similarly induced CPP (fig. S12, D and E). In contrast, the activation of male social memory engram did not induce conditioned place aversion, as observed in male subjects (Fig. 3K), likely due to variability across individuals. These findings indicate that sexual motivation toward opposite-sex conspecifics is not the sole determinant of the valence of the social memory engram.

To ascertain whether a female social memory engram specifically represents a female mouse rather than merely encoding general positive valence, we leveraged the propensity of male mice to reduce ultrasonic vocalizations (USVs) toward familiar female mice (*14*). We labeled a female social memory engram with ChR2 and optogenetically activated it at least 24 hours after labeling, during which the subject was presented with a familiar (engram-labeled) or a novel female mouse (fig. S13A). Simultaneously, we recorded the USVs emitted by the male subject mice (fig. S13, B and C). We observed that both the number and total duration of USVs were significantly reduced only when the engram was activated in the presence of the familiarized female (fig. S13, D and E), indicating that female memory engrams represent familiar female conspecifics per se rather than encoding general positive valence.

To determine whether vCA1 neurons are essential for CPP induced by live mice, we first performed a CPP test using female and male mice positioned opposite to each other (fig. S14A). We confirmed that female mice elicited both real-time and conditioned place preferences (fig. S14B). Next, CA1 pyramidal cell-specific Cre transgenic (Trpc4-Cre) mice (*5*) were injected with AAVs encoding Cre-inducible inhibitory opsin (AAV5-EF1α:DIO-eArch3.0-EYFP) or fluorescent protein (AAV5-EF1α:DIO-EYFP) into the bilateral vCA1 (fig. S14C), leading to robust transgene expression (fig. S14D). Optogenetic inhibition of vCA1 pyramidal cells during conditioning did not alter either real-time or conditioned place preferences (fig. S14, E and F), indicating that online neural activity in vCA1 is not essential for immediate induction or memory formation for place preference associated with live mice. In contrast, optogenetic inhibition of vCA1 pyramidal cells during the Post-test significantly suppressed the expression of CPP (fig. S14, G and H). To test the necessity of a specific social memory engram for expressing CPP, we labeled a female engram with an inhibitory opsin (NpHR), conditioned the subject with the labeled female, and inhibited the engram during the Post-test (Fig. 3, L and M). Engram inhibition significantly reduced CPP (Fig. 3, N and O). These findings indicate that neural activity in vCA1, specifically the social memory engram representing the female mouse, is essential for expressing CPP with live mice.

### dCA2 is necessary for high-dimensional encoding of social memory

The dorsal CA2 region of the hippocampus (dCA2) plays an essential role in encoding and recalling social memories (*2*, *15–17*) through projections to the downstream vCA1 (*4*, *17*). To investigate how dCA2 shapes social memory in vCA1, we genetically ablated dCA2 neurons by injecting AAVs encoding a Cre-inducible diphtheria toxin receptor (DTR; AAV2-CAG:FLEX-DTR-GFP) into the bilateral dCA2 of Map3k15-Cre mice that specifically target CA2 neurons (*18*) (Fig. 4A) and conducted activity-dependent double cell labeling (Fig. 4B). Intraperitoneal administration of diphtheria toxin (DT) 2 weeks after virus injection induced a significant loss of PCP4-positive cells in dCA2 (fig. S15). During the same surgery, AAV2-c-fos:tTA and AAV5-TRE:HTG were bilaterally injected into the vCA1 for activity-dependent double labeling (Fig. 4A). Unlike mice with intact dCA2 (Fig. 3E), ablation resulted in a similar number of HTG^+^c-Fos^+^ double-positive neurons and an unchanged proportion of double-positive neurons among HTG^+^ neurons across all three groups (Fig. 4, C and D). To assess the contribution of dCA2 to the sex-specific valence of social memory, we ablated dCA2 neurons and performed an optoCPP test (Fig. 4, E to G). Although dCA2 ablation did not diminish the positive reinforcing effect of activating female social memory, it did shift the valence of male social memory from negative to positive, leading to CPP (Fig. 4, H and I).

**Fig. 4.**
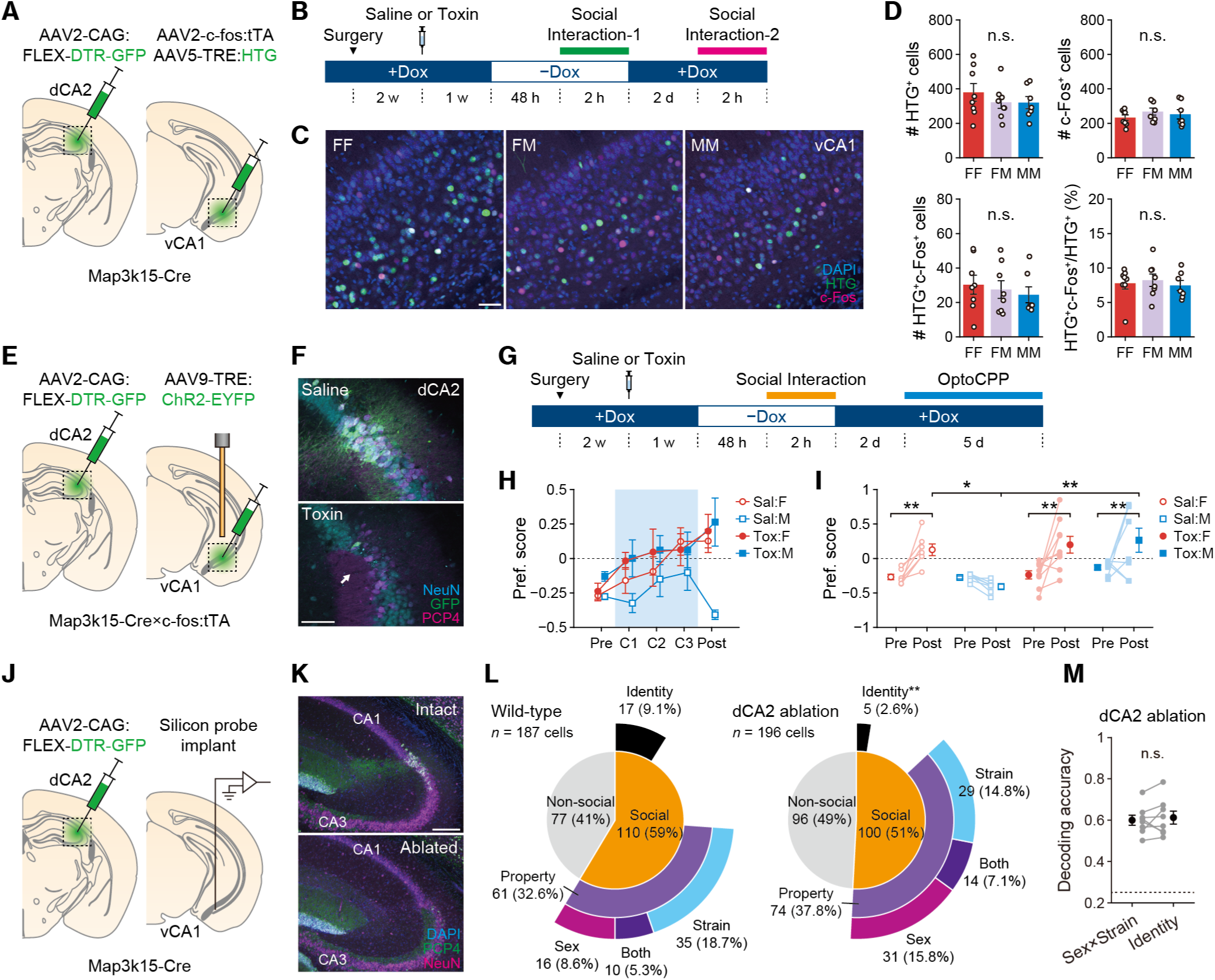
dCA2 is necessary for the high-dimensional encoding of social memory. (**A**) Schematic representation of dCA2 ablation using Cre-inducible diphtheria toxin receptor (DTR) and activity-dependent cell labeling. (**B**) Timeline of the double labeling of activated neurons after dCA2 ablation. (**C**) Representative images of the vCA1. Scale bar, 50 µm. (**D**) Left to right: number of HTG^+^ cells, number of c-Fos^+^ cells, number of HTG^+^c-Fos^+^ double-positive cells, and ratio of double-positive cells to HTG^+^ cells (*n* = 7–8 mice per group). (**E**) Schematic representation of dCA2 ablation using Cre-inducible DTR and activity-dependent cell labeling targeting. (**F**) Representative images of dCA2. Scale bar, 100 µm. (**G**) Timeline of the optoCPP test following dCA2 ablation. (**H**) Temporal dynamics of preference scores during conditioning (*n* = 8–9 mice per group). (**I**) Comparison of preference scores between Pre-test and Post-test. (**J**) Schematic representation of the dCA2 ablation and neural recordings. (**K**) Representative images of dCA2 after ablation. Scale bar, 200 µm. (**L**) The proportion of cell groups from wild-type (*n* = 187 cells from 6 mice across 6 sessions) and dCA2 ablated mice (*n* = 196 cells from 3 mice across 8 sessions). (**M**) Decoding accuracy of identity classifiers compared to the linear combination of decoding accuracy of sex and strain classifiers after dCA2 ablation. \**P* < 0.05, \**P* < 0.01, \*\*\**P* < 0.001; n.s., not significant. Error bars represent SEM.

To elucidate the circuit mechanism underlying the loss of population coding for social memory in vCA1, we recorded vCA1 neural activity using high-density silicon probes during the four-mouse social interaction test following dCA2 ablation (Fig. 4, J and K). Cell classification based on two-way ANOVA revealed a significant decrease in the proportion of cells with nonlinear interaction terms, including social identity cells (*P* = 0.0014, chi-square test of independence with post-hoc residual analysis; Fig. 4L). Consistently, the identity decoding accuracy became comparable to a linear combination of the sex decoding accuracy and strain decoding accuracies, reflecting the loss of nonlinear representation at the population level (Fig. 4M). These findings suggest that population coding of conspecific sex in social memory is relies on upstream dCA2, and its loss leads to a nondiscriminative positive valence of social memory regardless of sex.

### MeA is essential for the sex-specific valence of social memory

Neurons in the medial amygdala (MeA) exhibit sexually dimorphic responses to social stimuli (*19*, *20*), enabling pathway-specific control of distinct social behaviors (*21*, *22*). To investigate the role of the MeA in shaping sex-specific representations in social memory, we genetically ablated MeA neurons by injecting AAV2-CAG:FLEX-DTR-GFP and AAV2-EF1α:mCherry-IRES-Cre into the bilateral MeA of c-fos:tTA mice (Fig. 5A). Intraperitoneal administration of DT five weeks after virus injection caused a significant loss of GFP-positive neurons in the MeA (Fig. 5B and fig. S16). In line with previous studies (*23*, *24*), mice showed reduced female preference and mild deficits in social recognition memory one week after MeA ablation (fig. S17, A to F). As in the dCA2 experiment, AAV5-TRE:HTG was bilaterally injected into the vCA1 of c-fos:tTA mice and conducted activity-dependent double cell labeling (Fig. 5, A to C). Similar to dCA2-ablated mice (Fig. 4D), ablation resulted in a comparable number of HTG^+^c-Fos^+^ double-positive neurons and an unchanged proportion of double-positive neurons among HTG^+^ neurons across groups (Fig. 5D). To assess the role of the MeA in the sex-specific valence of social memory, we conducted an optoCPP test after MeA ablation (Fig. 5E). As observed with dCA2 ablation, MeA ablation induced CPP when the male social memory was labeled and reactivated (Fig. 5, F and G). Additionally, live male mice elicited CPP in male subject mice with MeA ablation (fig. S17, G and H). These results suggest that MeA ablation disrupts the representation of sex and sexual dichotomy of social memory valence.

**Fig. 5.**
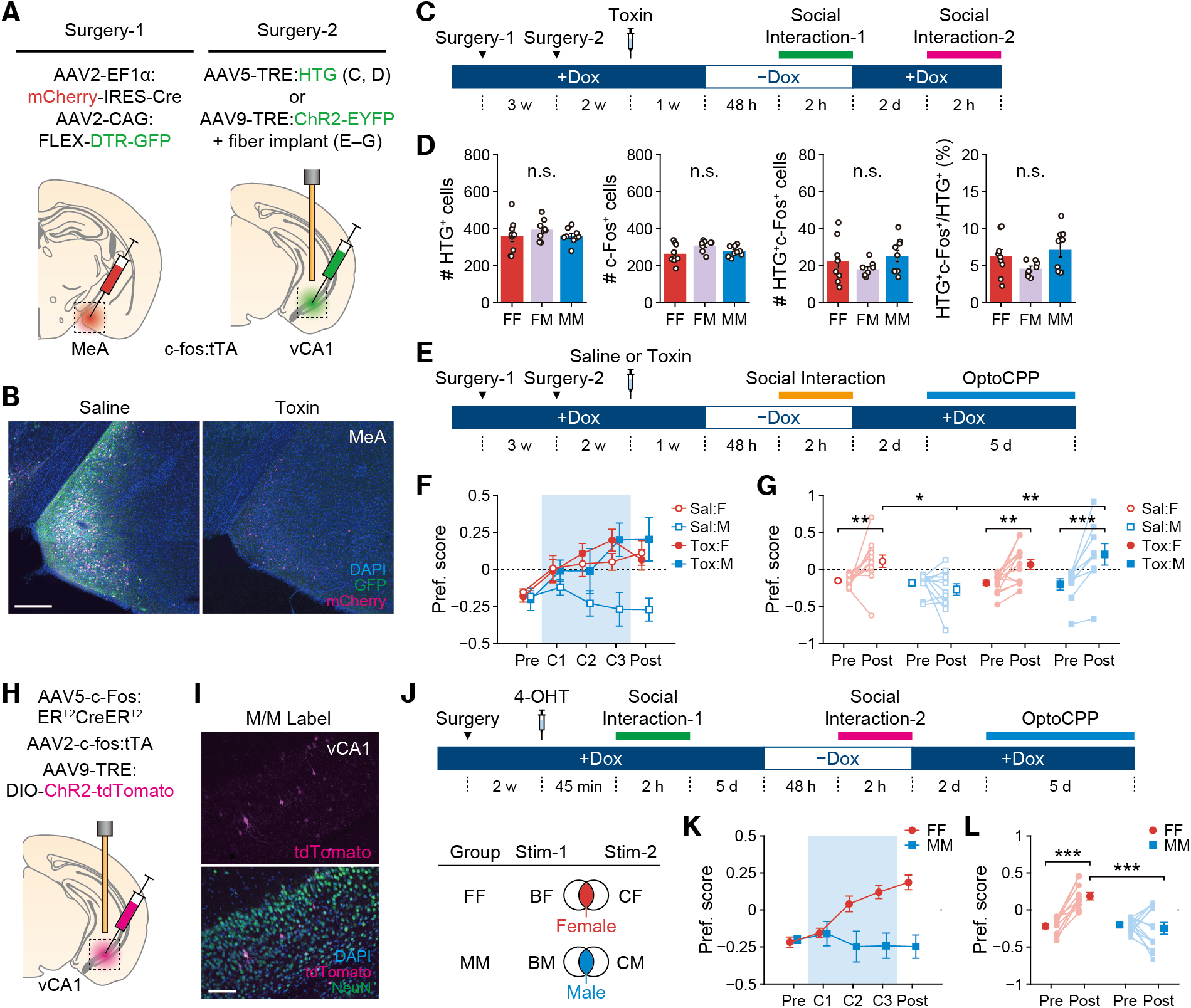
MeA is necessary for the representation of sex in social memory. (**A**) Schematic representation of MeA ablation using Cre-inducible diphtheria toxin receptor (DTR) and activity-dependent cell labeling. (**B**) Representative image of MeA. Scale bar, 200 µm. (**C**) Timeline of the double labeling of activated neurons after MeA ablation. (**D**) Left to right: number of HTG^+^ cells, number of c-Fos^+^ cells, number of HTG^+^c-Fos^+^ double-positive cells, and ratio of double-positive cells to HTG^+^ cells. (**E**) Timeline of the optoCPP test following MeA ablation. (**F**) Temporal dynamics of preference scores over conditioning (*n* = 9–13 mice per group). (**G**) Comparison of preference scores between Pre-test and Post-test. (**H**) Schematic representation of virus-mediated activity-dependent intersectional cell labeling with ChR2. (**I**) Representative images of the vCA1. Scale bar, 100 µm. (**J**) Experimental paradigm for optoCPP test following activity-dependent intersectional cell labeling with ChR2. Subject mice underwent two sessions of social interaction: first with a BALB/c female (BF) followed by a C3H female (CF) (FF group; *n* = 11 mice) or first with a BALB/c male (BM) followed by a C3H male (CM) (MM group; *n* = 13 mice). (**K**) Temporal dynamics of preference scores during conditioning. (**L**) Comparison of preference scores between Pre-test and Post-test. \**P* < 0.05, \*\**P* < 0.01, \*\*\**P* < 0.001; n.s., not significant. Error bars represent SEM.

### Intersectional activation of female social memories induces place preference

To ascertain whether the observed place preference is specifically associated with the activation of sexual representation in female social memory and whether such activation alone can elicit a positive reinforcing effect, we employed a two-step intersectional labeling technique. We injected AAVs encoding a TRE-inducible excitatory opsin (AAV9-TRE:DIO-ChR2-tdTomato), along with AAVs for an engineered Cre recombinase (AAV5-c-fos:ER^T2^CreER^T2^-PEST) and tTA (AAV2-c-fos:tTA), both of which are regulated by the *c-fos* promoter, into the bilateral vCA1 of wild-type mice (Fig. 5H). We confirmed that the c-fos:ER^T2^CreER^T2^ constructs function in an activity-dependent manner and are specific to the social stimuli (fig. S18). Initial social interactions in the presence of 4-hydroxytamoxifen (4-OHT) enabled *c-fos*-dependent Cre/lox recombination, which formed the basis for subsequent gene expression. Following a 1-week interval, to ensure the resetting of the CreER system, a second social interaction in the absence of Dox triggered *c-fos*-dependent activation of the tTA/TRE system that induced ChR2 expression (Fig. 5, I and J, and fig. S19). This sequence of manipulations ensured the exclusive expression of ChR2-tdTomato in neurons activated during both social interactions. Selective activation of neuronal populations shared between two female social memories, representing the female sex, leads to the development of place preference. In contrast, the activation of neurons associated with male representation did not produce this effect (Fig. 5, K and L).

## Discussion

Significant progress has been made in uncovering neural circuits that process sex and individual information in rodents, ranging from the vomeronasal system (*25*, *26*) to subcortical brain regions (*20*, *27*) and extending to higher-order cortical areas (*28*, *29*). In the present study, we demonstrated that mouse vCA1 neurons used two complementary encoding schemes to represent familiarized conspecifics. First, a subset of neurons displays nonlinear mixed selective responses (*30*, *31*) to a specific combination of sex and strain of individuals and are called social identity cells, which resemble concept-like cells in the human and primate hippocampus (*32*, *33*).

Second, a separate group of neurons responds specifically to either the sex or strain of individuals and collectively enables the construction of a low-dimensional, generalized representational map of social properties. The synergistic use of these two encoding strategies potentially facilitates an optimal population code for encoding familiarized conspecifics embedded in the social space (*34*) both in terms of robustness and efficiency (*35*, *36*).

The spike timing of hippocampal pyramidal cells undergoes a progressive shift from the ascending phase to the trough of hippocampal theta oscillations, a phenomenon known as phase precession (*37*, *38*), which collectively encodes the temporally compressed spatial sequence of an animal’s past, current, and future trajectories (*39*, *40*) as well as the nonspatial sequence of discrete events (*41*) or the passage of time (*42*). The hippocampal theta rhythm temporally organizes neuronal population activity in the vCA1 during social interactions (*6*), and the specific relationship between this organization and the representational geometry of social information is has not been elucidated. In this study, we demonstrated that neurons associated with social identity are predominantly activated around the trough of the theta rhythm, which is independent of social stimuli. In contrast, neurons that represent social properties tend to fire during the ascending phase and shift their preferences to an earlier phase during interactions with preferred stimuli. In the dorsal hippocampus, the spike timings of place cells are jointly influenced by two primary upstream afferent inputs at the theta timescale, one originating from the hippocampal CA3 region and the other from the entorhinal cortex layer 3 (EC3) (*43*), wherein a stronger EC3 input leads to a broader range of theta phase precession (*44*) particularly in a perceptual stimuli-enriched environment (*45*). Analogously, the spike timings of social property cells may be more strongly modulated by EC3 inputs than those of social identity cells, and leverage a wider range of hippocampal theta phases to construct a generalized map that represents multiple dimensions of social space.

The optogenetic reactivation of social memories linked to female, but not male, conspecifics exhibited a positive reinforcing effect. In addition, we demonstrated that intersectional reactivation of two distinct female social memories was sufficient to induce this effect. Consistent with the results of the electrophysiological recordings, these findings suggest that distinct neuronal populations in the vCA1 represent the sex information of familiar individuals. The valence of a social memory engram was determined by whether the stimulus was female or male, rather than by whether the stimulus was the same or opposite sex as the subject (Fig. 3K and fig. S12E). This sexual dichotomy in social memory valence may also stem from sexual dimorphism in neural circuits, including dCA2, as indicated by the female-specific involvement of dCA2 in contextual fear conditioning (*46*).

We demonstrated that ablation of the MeA, a key brain region essential for processing sexually dichotomous social sensory signals that emerge from the vomeronasal pathway (*20*, *25*, *26*), disrupts the representation of sex in the vCA1 and transforms the behavioral outcome of the activation of the social memory of male conspecifics into that of female conspecifics. Similarly, ablation of dCA2, which plays a pivotal role in encoding social memory (*2*, *3*), leads to the loss of high-dimensional population coding of familiar conspecifics (Fig. 4M) and nondiscriminative place preference by activating vCA1 social memories, irrespective of the sex of the stimuli (Fig. 4I). Activation of imperfect social memory, which lacks sex-specific information (fig. S20) or high-dimensional features of individual conspecifics, could exert a positive reinforcing effect by unmasking the inherently rewarding nature of social information (*47*), or by inducing an internal state akin to social novelty, which can elicit conditioned place preferences (*48*). Although functional connectivity is not fully understood, both the MeA and dCA2 send direct and indirect projections to the vCA1 (*4*, *49*, *50*). These afferents presumably entrain vCA1 social cells at specific theta phases with distinct frequencies that parallel CA3 and EC3 inputs, similar to place- and odor-encoding cells in the piriform cortex (*51*). Alternatively, the development of social property cells into social identity cells may represent a maturation process wherein, ultimately, multiple sensory inputs are integrated into a sparse conceptual representation of social identity (*33*, *52*).

## Acknowledgments

We thank T. Kitanishi and S. Fujisawa for their helpful comments on the manuscript revision, and all members of the Laboratory for Behavioral Neuroscience (Okuyama Lab) for their advice and support. We also thank The University of Tokyo IQB Olympus Bioimaging Center (TOBIC) for their assistance with microscopy and image acquisition.

## Author contributions

Conceptualization: AW, KT, TO; Methodology: AW, KT, TO; Investigation: AW, KT; Visualization: KT; Funding acquisition: AW, KT, TO; Supervision: TO; Writing – original draft: KT, TO; Writing – review & editing: AW, KT, TO

## Data and materials availability

All data are available in the main text and supplementary materials or have been deposited to Dryad (*53*).

## Supplementary Materials

Materials and Methods

Figs. S1 to S20

References (*1*–*53*)

