## Supplementary Material for "Representation of sex-specific social memory in ventral CA1 neurons"

**The PDF file includes:**

Materials and Methods
Figs. S1 to S20
References

### Materials and Methods

Mice

All procedures adhered to protocols approved by the Institutional Animal Care and Use Committee of the Institute for Quantitative Biosciences (IQB), University of Tokyo. All animals were socially housed under a 12-hour (7 a.m.–7 p.m.) light/dark cycle, with food and water *ad libitum*. C57BL/6J, C3H, and BALB/c mice were sourced from Central Laboratories for Experimental Animals (CLEA). c-Fos:tTA, Map3k15-Cre, and Trpc4-Cre transgenic mice were generated as described previously (*5*, *18*). All mouse lines carrying the *c-fos:tTA* transgene were raised on a 40-mg/kg doxycycline (Dox) diet until they underwent activated neuron labeling.

Viruses

The plasmid AAV5-TRE:HTG (pAAV-TRE:histone H2B-EGFP-2A-TVA receptor-2A-rabies glycoprotein) was purchased from Addgene and packaged by the University of Massachusetts Medical School (UMMS) with a titer of $1.4\times10^{13}$ vg/mL.

AAV5-EF1α:DIO-eArch3.0-EYFP, AAV5-EF1α:DIO-EYFP, AAV2-CAG:FLEX-DTR-GFP, and AAV9-EF1α:DIO-EYFP were purchased from Addgene, with a titer of $5.0\times10^{12}$, $3.3\times10^{12}$ $8.9\times10^{12}$, and $2.3\times10^{13}$ vg/mL, respectively. AAV5-c-Fos:$ER^{T2}CreER^{T2}$-PEST was purchased from UNC Vector Core, with a titer of $2.1\times10^{11}$ vg/mL. AAV2-EF1α:mCherry-IRES-Cre was packaged and purchased from ViGENE Bioscience, with a titer of $1.0\times10^{12}$ vg/mL.

AAV2-c-fos:tTA, AAV9-TRE:ChR2-EYFP, AAV9-TRE:NpHR-EYFP, AAV9-TRE:EYFP, AAV9-TRE:DIO-ChR2-tdTomato, and AAV9-TRE:DIO-tdTomato were generated in-house, with a titer of $2.6\times10^{11}$, $4.3\times10^{13}$, $1.1\times10^{11}$, $7.3\times10^{13}$, $1.3\times10^{12}$ , and $1.0\times10^{13}$ vg/mL, respectively. Specifically, the respective pAAV plasmids, along with the AAV helper plasmid (pAdDeltaF6; Addgene #112867) and either pAAV2/2 (for AAV2; Addgene#104963) or pAAV2/9n (for AAV9; Addgene #112865) were co-transfected into HEK-293T cells using PEI-Max for AAV production. Using the AAVpro Purification Kit Midi or Maxi (Takara Bio, Shiga, Japan), AAV was purified 3–5 days after transfection. The viral concentration was measured using qRT-PCR.

Surgery

**Silicon probe implant.** All surgical procedures for neural recordings were performed under 1–1.5% isoflurane anesthesia, and the mice were mounted on a stereotaxic apparatus (SR-9M-HT; Narishige). Dental cement (Super-Bond C&B, SUN MEDICAL) was used to attach a custom-made head frame onto the cleaned skull for the convenience of subsequent silicon-probe implantation. Ground and reference screws were implanted in the skull above the cerebellum. To ablate dCA2 neurons, Map3k15-Cre mice were injected with AAV, as described below, during the same surgery. The mice were housed individually postoperatively. The mice were allowed to interact with their littermates in their home cages for 5–10 minutes per day to maintain sociability. After 2–3 weeks of recovery, the mice were again mounted with the head frame, and 128-channel silicon probes (P128-5; Diagnostic Biochips), attached to movable microdrives, were implanted targeting the right ventral hippocampus (−2.80 to –3.25 mm AP, +3.20 mm ML from bregma) parallel to the midline, facing the channels laterally. To visualize the tracks, the probes were immersion-coated with CM-DiI (V22888; Invitrogen) dye solution before surgery.

**Virus injection and fiber implantation.** Mice were anesthetized with an intraperitoneal administration of an anesthetic mixture containing 0.75 mg/kg medetomidine (Domitor, Orion Corporation), 4 mg/kg midazolam (Sandoz), and 4 mg/kg butorphanol (Vetorphale, Meiji Animal Health) and mounted on a stereotaxic apparatus. Viruses were injected at a flow rate of 2 nL/s using a glass micropipette attached to a 10-µL Hamilton microsyringe, controlled using a micropump (UMP3, World Precision Instruments) and a controller. Injections were targeted bilaterally to the vCA1 (−3.16 mm AP, ±3.10 mm ML, −4.70 mm DV from Bregma), MeA (−1.58 mm AP, ±1.90 mm ML, −5.50 mm DV), and dCA2 (−1.60 mm AP, ±1.60 mm ML, −1.70 mm DV). The pipette was slowly lowered to the target site and left in situ for 5 minutes postinjection. The incision was closed using sutures. Postoperatively, mice were administered 5 mg/kg carprofen (Remadile, Zoetis) for postoperative analgesia and 0.75 mg/kg atipamezole (Antisedan, Orion Corporation) for the reversal of the sedative and analgesic effects of medetomidine and remained on a heating pad until they had fully recovered from the anesthesia.

For optoCPP experiments using the reactivation of social memory neurons, 150 nL of AAV9-TRE:ChR2-EYFP or AAV9-TRE:EYFP was injected bilaterally into the vCA1 of c-fos:tTA transgenic mice. For the optoCPP experiments using the intersectional reactivation of social memory neurons, 150 nL of a 1:1:8 cocktail of AAV5-c-Fos:$ER^{T2}CreER^{T2}$-PEST, AAV2-c-fos:tTA, and AAV9-TRE:DIO-ChR2-tdTomato was injected bilaterally into the vCA1 of C57BL/6J wild-type mice. For optogenetic whole inhibition experiments, 150 nL of AAV5-EF1α:DIO-eArch3.0-EYFP or AAV5-EF1α:DIO-EYFP was injected bilaterally into the vCA1 of Trpc4-Cre mice. For optogenetic engram inhibition experiments, 150 nL of AAV9-TRE:NpHR-EYFP or AAV9-TRE:EYFP was injected bilaterally into the vCA1 of c-fos:tTA transgenic mice. To validate the expression of the c-Fos:$ER^{T2}CreER^{T2}$ construct, 150 nL of a 1:1:8 cocktail of AAV5-c-Fos:$ER^{T2}CreER^{T2}$-PEST, AAV2-c-fos:tTA, and AAV9-TRE:DIO-tdTomato was injected bilaterally into the vCA1 of C57BL/6J wild-type mice. For USV recordings with female engram activation, 150 nL of a 1:4 cocktail of AAV2-c-fos:tTA and AAV9-TRE:ChR2-EYFP was injected bilaterally into the vCA1 of C57BL/6J wild-type mice. To label the activated neurons followed by c-Fos immunohistochemistry, 150 nL of a 1:4 cocktail containing AAV2-c-fos:tTA and AAV5-TRE:HTG was injected bilaterally into the vCA1 of C57BL/6J wild-type mice. To ablate MeA neurons, 150 nL of a 1:5 cocktail containing AAV2-EF1α:mCherry-IRES-Cre and AAV2-CAG:FLEX-DTR-EGFP was injected bilaterally into the MeA of c-fos:tTA transgenic mice. To ablate dCA2 pyramidal neurons, 150 nL of AAV2-CAG:FLEX-DTR-GFP was injected bilaterally into the dCA2 of Map3k15-Cre transgenic mice (for neural recordings during the four-mouse social interaction test) or Map3k15-Cre×c-fos:tTA double transgenic mice (for optoCPP tests).

For optogenetic manipulation experiments, following the viral injection during the same surgery, optic fibers (200-µm diameter, NA 0.22) were implanted by targeting to the bilateral vCA1 (−3.16 mm AP, ±3.10 mm ML, −4.5 mm DV from Bregma). For additional stability, two screws were inserted into the skull surrounding the implant site in each hemisphere. A layer of adhesive cement was applied, followed by dental cement, to secure the optical fiber implant. The implanted fibers were protected using a custom-made 3D-printed cap.

Cell ablation

After 5 weeks (for MeA) or 2 weeks (for dCA2) following the virus injection to induce the Cre-dependent expression of diphtheria toxin receptor (DTR), mice were administered 50 µg/kg of diphtheria toxin (DT) or saline via intraperitoneal injection. One week later, mice underwent double-labeling, optoCPP, or neural recordings.

Cell labeling

**Labeling with doxycycline.** Three weeks after the virus injection, the mice were given doxycycline (Dox)-free food for 48 hours to establish a labeling window. Subsequently, a female or male mouse of BALB/c or C3H strain was introduced into the subject mouse's home cage as a social stimulus to label the neurons activated by the stimulus. Following a 2-hour social interaction, the stimulus mouse was removed, and food containing 40 mg/kg of Dox was reinstated. Two days later, mice underwent optoCPP, USV recording, or double-labeling experiments.

**Labeling with 4-hydroxytamoxifen.** 4-hydroxytamoxifen (4-OHT; H6278, Sigma-Aldrich) was dissolved in castor and sunflower oils. Mice were intraperitoneally injected with 50 mg/kg of 4-OHT or the oil solvent alone (the oil control group for validating the expression of c-Fos:$ER^{T2}$Cre$ER^{T2}$ construct). Forty-five minutes later, the mice underwent 2 hours of social interaction in their home cages.

**Labeling with c-Fos protein expression.** For the double-labeling of activated neurons, a female or male C3H mouse was introduced into the subject mouse's home cage for the second labeling. Following a 2-hour interaction, the stimulus mouse was removed, and the subject mouse was perfused for subsequent immunohistochemistry and cell counting.

Behavior

**Four-mouse social interaction test.** The mice were acclimatized to the social arena and social chambers for 5 minutes daily for 2–3 days. The social arena was a white acrylic square box (38×38 cm area, 30 cm height), and two custom-made social chambers with quadrant bases (7.5-cm radius; two of six subjects) or circular bases (8-cm diameter; four of six subjects) were 3D printed (Original Prusa i3 MK3S, Prusa Research) using white PETG filaments placed at opposite corners of the arena. One day prior to the test, the subject mice were placed in a familiarization chamber along with four stimulus mice consisting of female and male BALB/c and C3H mice. The familiarization chamber was a white acrylic rectangular box (24.5×31 cm area, 25 cm height) composed of one large central room (24.5×12 cm) where the subject mice were placed and four smaller rooms (12×8 cm) where each stimulus mouse was positioned. Large and small rooms were separated using a stainless-steel wire mesh with ~1 cm gaps.

On the test day, after indirect familiarization with the stimulus mice for approximately 24 h, the subject mice and each stimulus mouse were allowed to interact directly with each other for 10 minutes in the subject mouse's home cage. Following direct habituation, mice were subjected to five consecutive trials. After the first trial, which took place in the absence of social stimuli (E-E trial), two stimulus mice were placed in the left or right chamber, and the subject mouse was allowed to explore the arena for 5 minutes (CM-CF trial). This procedure was repeated by alternating the two stimulus mice with the three other combinations (BF-BM, BM-CM, and CF-BF trials). Trials were conducted at 5-minute intertrial intervals. Trials were video recorded at 25 frames per second from a top view of the arena using an area scan camera (acA1300-60gmNIR; Basler) fitted with a Computar 4–8-mm C-Mount lens. The video recording system was controlled using the Bonsai software (*54*).

**Optogenetic conditioned place preference test.** The test chamber was a white acrylic rectangular box (20×48.6 cm area, 25-cm height) composed of two side zones (20×20 cm) and a central zone (20×8 cm) that was connected to the side zones via 8-cm openings. The left zone featured a black-striped wall and square-punched floor, whereas the right zone featured a black-dotted wall and round-punched floor. The locations of the subjects were continuously tracked and recorded in real time using the Noldus EthoVision XT software.

The optogenetic conditioned place preference (optoCPP) test consisted of three phases over 5 days. On Day 1, the mice were placed in the central zone and allowed to freely explore the entire chamber for 10 minutes (pre-test). The zone for which the mice showed less preference during the pretest was identified as the target zone for light stimulation. On days 2–4, the mice were again placed in the chamber for 10 minutes, and light stimulation was delivered while the mouse was in the target zone (conditioning). The entrance to the target zone was detected using the EthoVision software, triggering a stimulus generator (STG-4008, Multichannel Systems) through a Noldus USB-IO Box. The stimulus generator drove a 473-nm laser (CNI Laser) to deliver 20-Hz light stimulation with a duty cycle of 30%. The laser output was tested at the start of each experiment to confirm the delivery of at least 10 mW of power to the ends of the optical fiber patch cords. On Day 5, similar to Day 1, the mice were placed in the central zone and allowed to freely explore the entire chamber for 10 minutes (post-test).

**Conditioned place preference test using social stimuli.** The CPP test using live mice as stimuli was conducted in the same test chamber as that for the optoCPP test described above, with the modification that a female and/or male C3H mouse was presented during the conditioning phase instead of labeling and reactivating the social memory neurons in the vCA1. Social stimuli were confined to custom-made social chambers featuring circular bases (8 cm diameter) positioned at the top-left corner of the left zone and the bottom-right corner of the right zone. For tests with a pair of female and male mice as stimuli, female mice were assigned to the zone where the subject showed less preference during the pre-test, while male mice were assigned to the opposite zone. For tests using a male mouse as stimuli, it was assigned to the zone were the subject showed less preference during the pre-test, whereas the opposite social chamber was kept empty.

Optogenetic whole inhibition experiments were conducted 2 weeks after virus injection. The subject mice were connected to optic fiber patch cords in each trial for 5 days. During the conditioning (days 2–4) or the post-test (day 5), 561-nm laser light with 15-mW power was delivered while the mouse was in the target zone.

Optogenetic engram inhibition experiments were conducted 3 weeks after virus injection. Subject mice were connected to optic fiber patch cords in each trial for 5 days. During the post-test (day 5), 589-nm laser light with 15-mW power was delivered while the mouse was in the target zone.

**Sexual preference test.** A rounded rectangular arena was used in the sexual preference test. This arena comprised a 30×30 cm square and, on either side, two half-circles with a 15-cm radius and a height of 30 cm. One week after receiving intraperitoneal injections of either saline or DTR for MeA ablation, mice were allowed to explore the arena for 10 min. During the test, female and male C3H mice were confined to pencil holders (each 7.5 cm in diameter and 15 cm in height) and positioned in the center of the half-circles of the arena. Entrance into the social zone (defined as 1.5 times the diameter of the pencil holder) was monitored, recorded, and analyzed using EthoVision software.

**Social recognition memory test.** After the sexual preference test, mice underwent a social recognition memory test. In the same arena, two novel social stimuli (male BALB/c and male C3H mice) were confined to a pencil holder and positioned in the arena. The mice were initially allowed a 10-minute exploration (Pre-test) and subsequently returned to their home cages, after which the C3H male mouse was introduced into the subject's home cage for 2 hours. Following the removal of the familiarized mouse and a 30-minute interval, the subjects were again placed in the arena, now containing the same BALB/c and C3H mice, for a 10-minute exploration (Post-test). The entrance into the social zone, as in the previous test, was monitored, recorded, and analyzed using EthoVision software.

**Social recognition memory test with female engram activation.** Sexually experienced C57BL/6J wild-type mice were injected with an AAV cocktail for *c-fos*-dependent ChR2 expression as described above and implanted with optic fibers targeting the bilateral vCA1. Three weeks after surgery, social memory engrams were labeled through a 2-hour social interaction with a female C3H mouse. Two days after labeling, the subjects underwent one 3-minute trial per day for four consecutive days. During each trial, the subject mice were connected to optic fiber patch cords and allowed to explore the social arena in the presence of a freely moving female mouse. The social arena consisted of a white acrylic square box identical to that used in the four-mouse social interaction test. The trials followed a 2×2 design, with the factors being the type of social stimulus (a familiar C3H or a novel C57BL/6J) and the presence or absence of light stimulation (473-nm, 20 Hz, 30% duty cycle; identical to those used in the optoCPP tests but applied throughout the trial). The order in which each combination of conditions (social and light stimulation) was presented was counterbalanced across the four trials and subjects.

USV recording and analysis

Ultrasonic sounds were recorded using a condenser ultrasound microphone (CM16/CMPA, Avisoft Bioacoustics), and audio signals were digitized at 375 kHz with an ultrasound recording interface (UltraSoundGate 116H, Avisoft Bioacoustics). USV syllables were detected using the USVSEG software ver 0.9 (rev 2) (*55*). Syllables were defined as having a frequency range of 40–160 kHz, a duration of 3–300 ms, and an amplitude greater than 4.5 standard deviations (SD).

Histology

At the end of the experiment, the animals were deeply anesthetized with isoflurane, and transcardially perfused with phosphate-buffered saline (PBS) followed by 4% paraformaldehyde (PFA) in PBS. Brains were extracted, postfixed in 4% PFA for 24 h, and then sliced at 50- or 100-µm thickness (only after the neural recordings) using a vibratome (LEICA VT1000 S). Images were acquired using a fluorescence microscope (Keyence BZ-X710) or a confocal microscope (Evident FV3000).

For immunohistochemistry, brain sections were blocked with 5% normal goat serum (NGS) in PBST (PBS with 0.3% Triton-X100) for 1 hour at room temperature (RT). The sections were incubated with primary antibodies in PBST containing 5% NGS for 24 h at 4°C. The primary antibodies used were chicken anti-GFP (1:1000; Invitrogen), mouse anti-NeuN (1:1000; Merck), guinea pig anti-c-Fos (1:1000; Synaptic Systems), rabbit anti-PCP4 (1:250, Merck), and rabbit anti-RFP (1:1000; ROCKLAND). After washing three times with PBS for 15 minutes each, the sections were incubated with secondary antibodies conjugated to Alexa488, Alexa 555, or Alexa 647 (1:500; Invitrogen) for 3 hours at RT. The sections were washed three times with PBS for 15 minutes each, and DAPI (1:1000) staining was performed during the second wash. Sections were mounted using Fluoromount/Plus (Diagnostic BioSystems) and imaged using a fluorescence microscope.

Cell counting

Coronal sections containing vCA1 were prepared from the brains of mice following the cell labeling experiment. Immunohistochemical staining was performed as previously described. Fluorescence images from a single optical plane were acquired using a microscope equipped with a 10× or 20× objective lens and subsequently analyzed using ImageJ software. A threshold of mean + 2 SD over the entire pixel range of the image was applied for each color channel, and positive cells were identified through manual inspection. The number of cells in each ROI was normalized to the size of the standard ROI.

Neural recordings

The probes were connected to the Open Ephys data acquisition system (*56*) and controlled using the Open Ephys GUI. The probes were gradually lowered over several days until the CA1 pyramidal layer was reached, as determined by the appearance of hippocampal SPW-Rs and pyramidal cell activity. Electrophysiological data were sampled at 30 kHz.

Spike sorting and unit classification

Spike sorting was performed automatically using Kilosort2.5 (*57*). This was followed by manual adjustment of the waveform clusters using the Phy2 software (*58*). Units with firing rates lower than 0.1 Hz were omitted. Units were classified as putative pyramidal cells if the overall firing rate was < 15 spikes/s, the trough-to-peak length of spike shapes was > 0.425 ms, and the exponential time constant of the rise time in an autocorrelogram ($\tau_{rise}$) was < 10 ms.

LFP analysis

The wideband neural signal was downsampled to 1.25 kHz and used as local field potential (LFP) signal. The channel with the highest theta power in the normalized spectrum, located in the middle to deep sublayers of the CA1 pyramidal cell layer, was selected for the theta cycle detection. Theta-band signals were extracted by applying a bandpass filter (6–12 Hz) to the LFP signals. Hilbert transformations were then employed to estimate the instantaneous theta phase, where a 0-degree phase corresponds to the peak and a ±180-degree phase to the trough.

Behavioral analysis

For the neural recording experiments, the positions of the noses of the subject mice were detected using SLEAP software (*59*). Animal trajectories were smoothed using a 5-bin sliding window. The relative social distance (*d*) between the subject mouse and the two stimulator mice in each frame was calculated as $(d_L - d_R)/(d_L + d_R)$, where $d_L$ and $d_R$ represent the distances between the subject's nose position and the respective centers of the left and right social chambers where the stimulus mice were located. Social interaction events were defined as periods during which the relative distances had $|d| \geq 0.72$ (for quadrant chambers) or $|d| \geq 0.81$ (for circular chambers), which closely corresponded to the boundaries of the social chamber. The precise discrepancy between the boundary of the social chamber and the social zone defined by the relative distance (at most 0.66 cm) was negligible when considering the spatial resolution of the video data (0.06 cm/pixel) and the tracking accuracy of our trained SLEAP model (prediction error for the 95$^{th}$ percentile: 8.2 pixels). Events with < 0.5 second time intervals were consolidated, whereas those lasting < 1 second were eliminated.

For the conditioned place preference test, the preference score was calculated as $(t_P - t_N)/(t_P + t_N)$ where $t_P$ and $t_N$ are the length of time that the subject mouse stayed in the zone where the stimulus was placed (P) and not placed (N). Analogously, the preference score for the social recognition memory test was calculated as $(t_N - t_F)/(t_N + t_F)$ where $t_N$ and $t_F$ are the length of time that the subject mouse interacted with the nonfamiliarized mouse (N) and the familiarized mouse (F), respectively.

Firing rate maps

For visualization purposes, firing rate maps were obtained as follows. First, spike timings were assigned to the corresponding nose positions of the subjects and arranged into 1cm×1cm spatial bins. The number of spikes for each bin was then divided by the bin occupancy time. The maps were smoothed using a 5×5 box window filter (*60*) with a weight of:

```
w = [ ···
  0.0025 0.0125 0.0200 0.0125 0.0025; ···
  0.0125 0.0624 0.1000 0.0625 0.0125; ···
  0.0200 0.1000 0.1600 0.1000 0.0200; ···
  0.0125 0.0625 0.1000 0.0625 0.0125; ···
  0.0025 0.0125 0.0200 0.0125 0.0025]
```

Single-cell classification analysis

Two-way ANOVA was performed to explore the influence of sex and strain of social stimuli on the z-scored firing rates of individual neurons during social interactions. This analysis incorporated the main effects of sex and strain along with their interaction effects. Statistical significance was set at a threshold of $P < 0.05$. Neurons exhibiting significant main effects of sex or strain or a significant interaction effect were classified as social neurons. In contrast, neurons that did not show significant main or interaction effects were classified as non-social neurons. Additionally, a chi-square goodness-of-fit test was used to evaluate the homogeneity of subpopulations within the sex or strain categories of the neurons.

For visualization purposes, the differences in the mean firing rates across social interaction events, grouped by the properties of the social stimuli were computed.

Cell pair analysis

The significance of theta modulation in spike cross-correlograms (CCGs) was assessed by calculating the power spectrum density (PSD) from both the real and permuted spike timings of neuron pairs. Spike timings were extracted for each neuron during social interactions, and CCGs were computed for each neuron pair (bin size: 1 ms; total width: ±1 s). PSDs were calculated using Welch's method, focusing on the area under the curve (AUC) within the theta frequency band (6–12 Hz). For the construction of null models, spike timings were circularly shifted across the entire session and theta AUCs were recalculated. This procedure was repeated 1,000 times to determine statistical significance. The *P*-values obtained from the permutation test were converted to z-scores using the following equation:

$$Z = \Phi^{-1}\left(\left|\frac{P}{2}\right|\right)$$

where $\Phi^{-1}$ is the inverse of the standard normal cumulative distribution function.

To detect temporal shifts in the CCG peak, CCGs were smoothed using a Gaussian kernel (standard deviation: 12.5 ms), and the absolute difference between the zero point and the first peak within a ±125-ms window (equivalent to a cycle of an 8-Hz rhythm) was measured as the shift value. The pairwise distance in neuronal activity (*D*) between two cells was determined using the cosine similarity, expressed as

$$D = 1 - \frac{\sum_{i=1}^{N} A_i B_i}{\sqrt{\sum_{i=1}^{N} A_i^2} \cdot \sqrt{\sum_{i=1}^{N} B_i^2}}$$

where $A_i$ and $B_i$ represent the z-scored firing rates of neurons A and B, respectively, during the $i$-th $N$ social interaction. To correct for bias due to variations in trial numbers across subjects, a subset of trials equal to the minimum observed number ($n$ = 62) was randomly selected and used to compute the pairwise distances. This process was repeated 100 times to obtain average distances.

Population decoding analysis

Decoding was performed using linear support vector machine (SVM) classifiers with five-fold cross-validation implemented using the *fitcecoc* function in MATLAB. The firing rates of simultaneously recorded putative pyramidal cells during each social interaction were used as predictors. Identity decoding uses a one-versus-one approach. Subsampling for five-fold cross-validation was stratified according to sex and strain labels. The cross-validation was repeated 100 times, and the overall decoding accuracy was recorded as the mean across 100 repetitions. For decoder time courses around nose-poke onsets, 200-ms non-overlapping time bins were used to construct the decoders. To evaluate the decoding performance, a null model was generated by circularly shifting the event labels using random integers, and cross-validation was performed five times. This process was repeated 100 times, resulting in an overall pseudo-decoding accuracy that was determined from an average of 500 repetitions.

The weights assigned by the linear classifiers to each neuron are interpreted as indicators of relative importance. For the intersession comparison, the weights were normalized as follows:

$$\widehat{w}_i = \frac{N}{\sum_{k=1}^{N} |w_k|} \cdot |w_i|$$

where $w_i$ is the decoder weight of the $i$-th neuron, the hat symbol (^) indicates normalization, and $N$ is the total number of pyramidal cells recorded in each session. The relative importance of the 500 pseudo-decoders was calculated to assess the significance of the correlation between sex and strain.

Pseudo-population analysis

Pseudo-population activity matrices were constructed by concatenating the z-scored neuronal activity matrices from each mouse. As different sessions contained varying numbers of social interactions, a random subset of interaction bouts was excluded. This ensured that the number of bouts of each social stimulus was consistent and equal to the minimum number of bouts observed across all mice. Consequently, the size of the pseudo-population activity matrix is defined as follows:

$$\left[\sum_{i=1}^{N} n_i \, , \sum_{s\in S} \min_i\{\#I_i^s\}\right]$$

where $n_i$ is the number of recorded pyramidal cells from the $i$-th subject ($n$ = 15–57 cells), $N$ is the number of the subjects ($N$ = 6 mice), and $\#I_i^s$ represents the number of interaction bouts with the social stimulus $s \in \{BF, BM, CF, CM\}$ for the $i$-th subject.

Cross-condition generalization performance (CCGP)

To calculate CCGP (*10*), the interaction bouts were categorized based on the properties of the social stimuli. Specifically, the training set was composed exclusively of bouts from one set of properties, whereas the test set included only bouts from a different nonoverlapping set of properties. For instance, to assess the cross-sex strain CCGP, we trained a linear classifier using interaction data involving female BALB (BF) and C3H (CF) mice. Subsequently, through interactions involving male BALB (BM) versus C3H (CM) mice, we evaluated the ability of the classifier to distinguish between strains. This process was then inverted with the training and test datasets swapped, and CCGP was calculated as the mean decoding accuracy across both directions.

This procedure was repeated 100 times using randomly generated pseudo-population activity matrices to evaluate CCGP in the actual data. For each pseudo-population activity matrix, we estimated the CCGP of a null model by randomly shuffling the activity vectors of each neuron 100 times, resulting in 10,000 sets of CCGP values based on the permuted data. The significance of the real CCGP was determined by comparing it with the distribution of the CCGPs derived from the permuted datasets.

Spike-LFP analysis

Spike spectrograms, normalized power of LFP, and spike-LFP coherograms around social interactions were computed using the *cohgramcpt* function of the Chronux toolbox (http://chronux.org/). The significance of spike-LFP coherograms was determined using a permutation test.

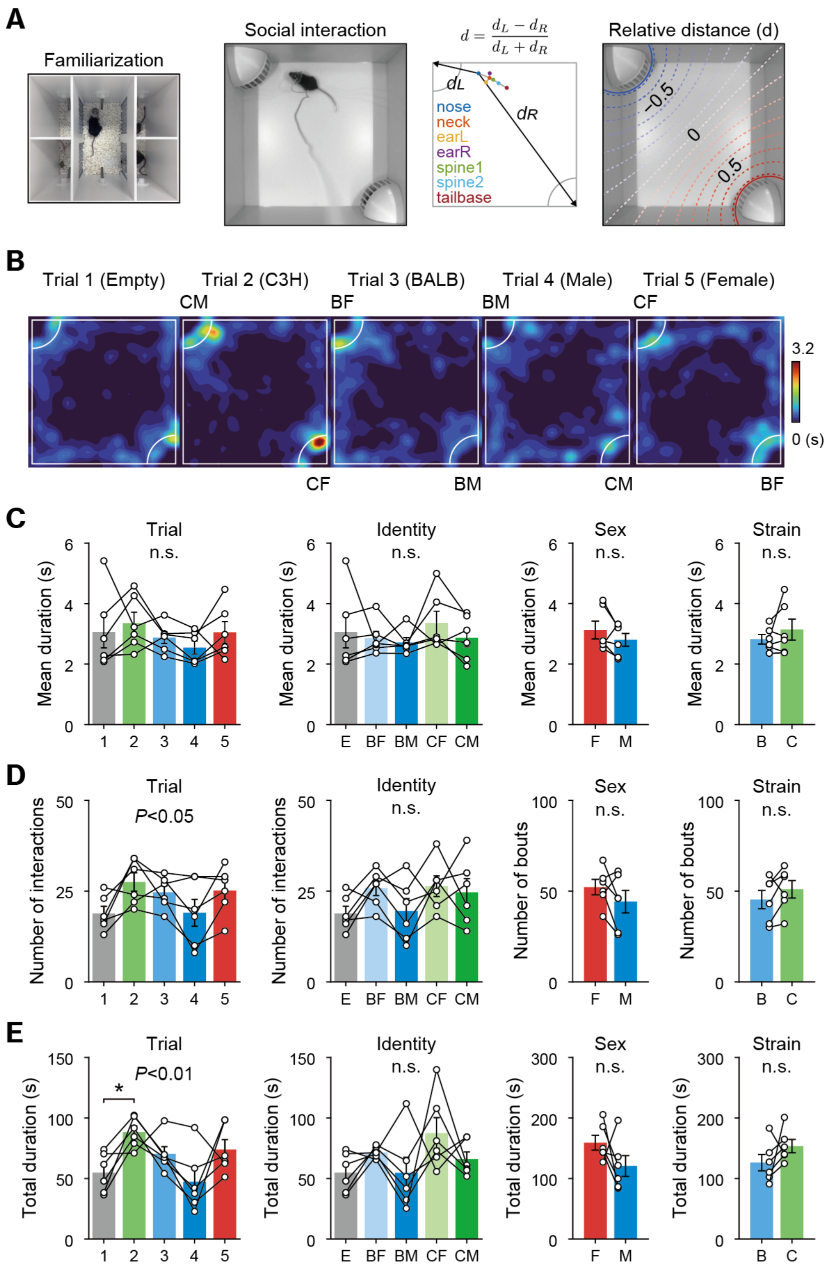


**fig. S1. Behavioral analysis during the four-mouse social interaction (FMSI) test.**

(**A**) From left to right: top view of the isolation chamber used for familiarization with BALB/c and C3H female and male mice, the arena used for social interaction, the definition of relative distance, and the relative distance projected onto the arena. (**B**) Representative position heatmaps from the FMSI test. (**C**–**E**) Mean bout duration (C), number of interaction bouts (D), and total bout duration (E) of social interaction. Significance was determined by one-way ANOVA (for Identity) or paired *t* test (for Sex and Strain).

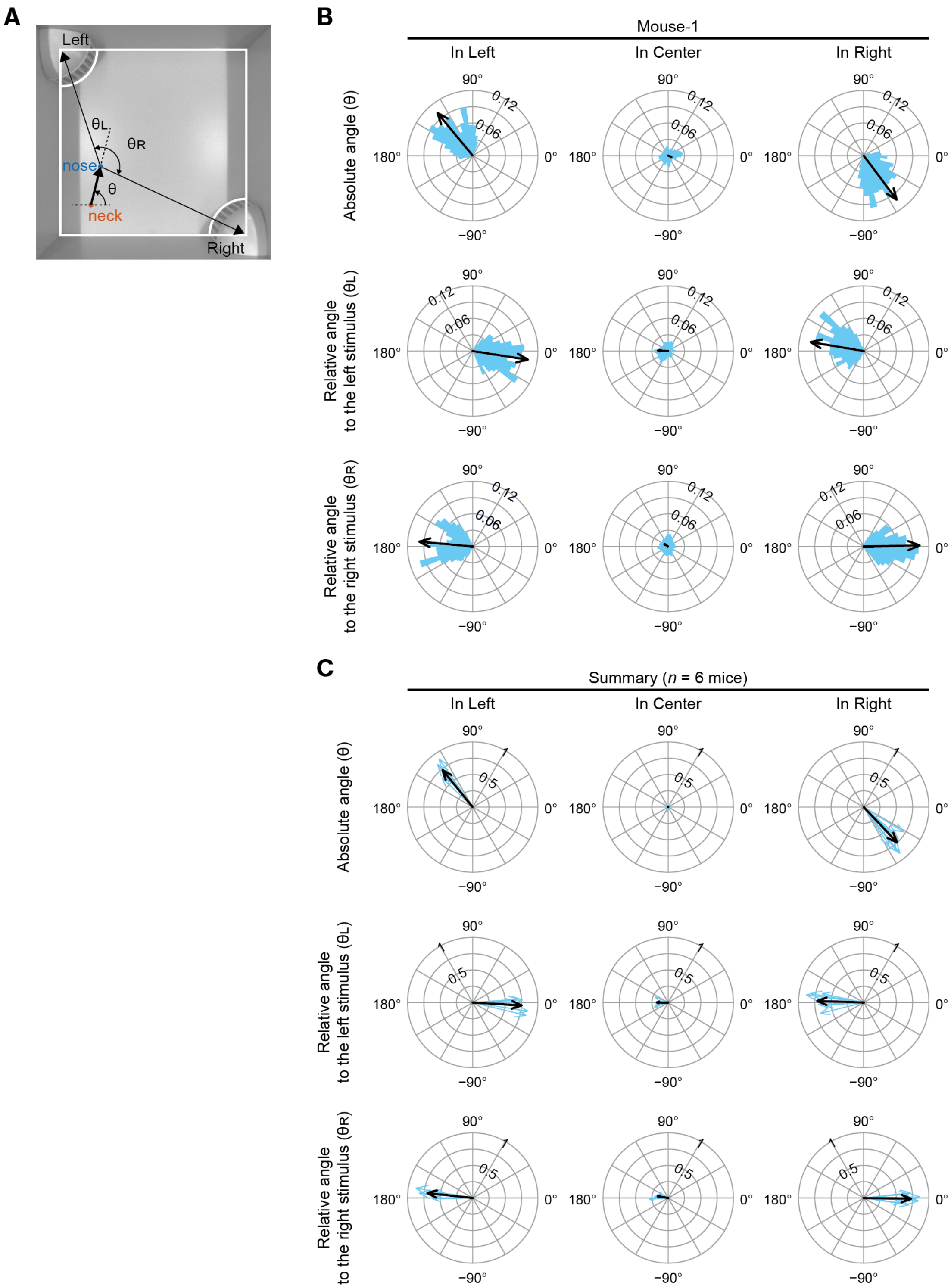


**fig. S2. Head direction analysis during the FMSI test.**

(**A**) Definition of head direction in the FMSI test. $\theta$ represents the absolute head direction, whereas $\theta_L$ and $\theta_R$ represent the relative head directions toward the left and right social chambers, respectively. (**B**) Distribution (shown as a polar histogram in blue) and circular mean (depicted by a black arrow) of the absolute head direction (top), relative head direction toward the left (middle), and right (bottom) social chambers of an example mouse. (**C**) Summary of the absolute and relative head direction ($n$ = 6 mice). Blue arrows indicate individual mice, and black arrows indicate the mean direction across subjects.

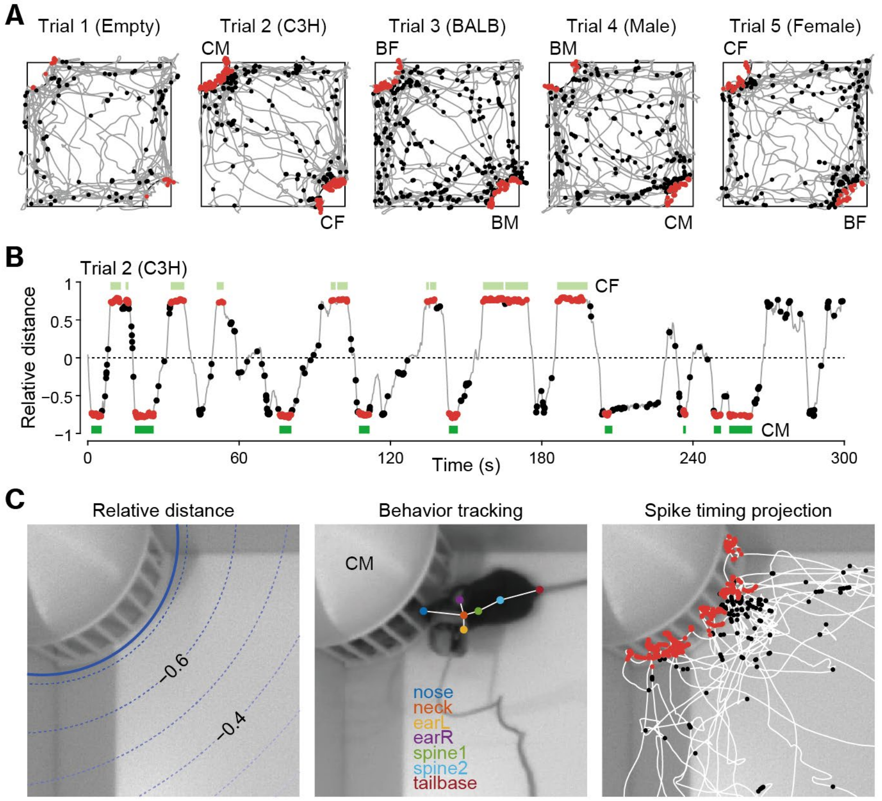


**fig. S3. Spatial dynamics of neuronal spike timing during social interaction.**

(**A**) Spike timings of an example cell projected onto behavioral trajectories for each trial. The red and black dots indicate spike timings that occurred during and outside social interactions, respectively. (**B**) Spike timings of the same cell projected onto the relative distance during trial 2. Light and dark green horizontal bars indicate periods of social interaction with the CF (right chamber) and CM (left chamber), respectively. (**C**) Magnified view of the left social chamber. Left: The threshold of relative distance (±0.72) corresponds to the social chamber's border. Center: The moment when the subject mouse inserted its nose into the social chamber, with post-hoc detected body points and skeleton overlaid. Right: Same as (A) but overlaid on the top view of the social arena. Note that some spikes observed outside the social zone were included in the social interaction period because two adjacent interaction events separated by transient (< 0.5 s) exits were concatenated.

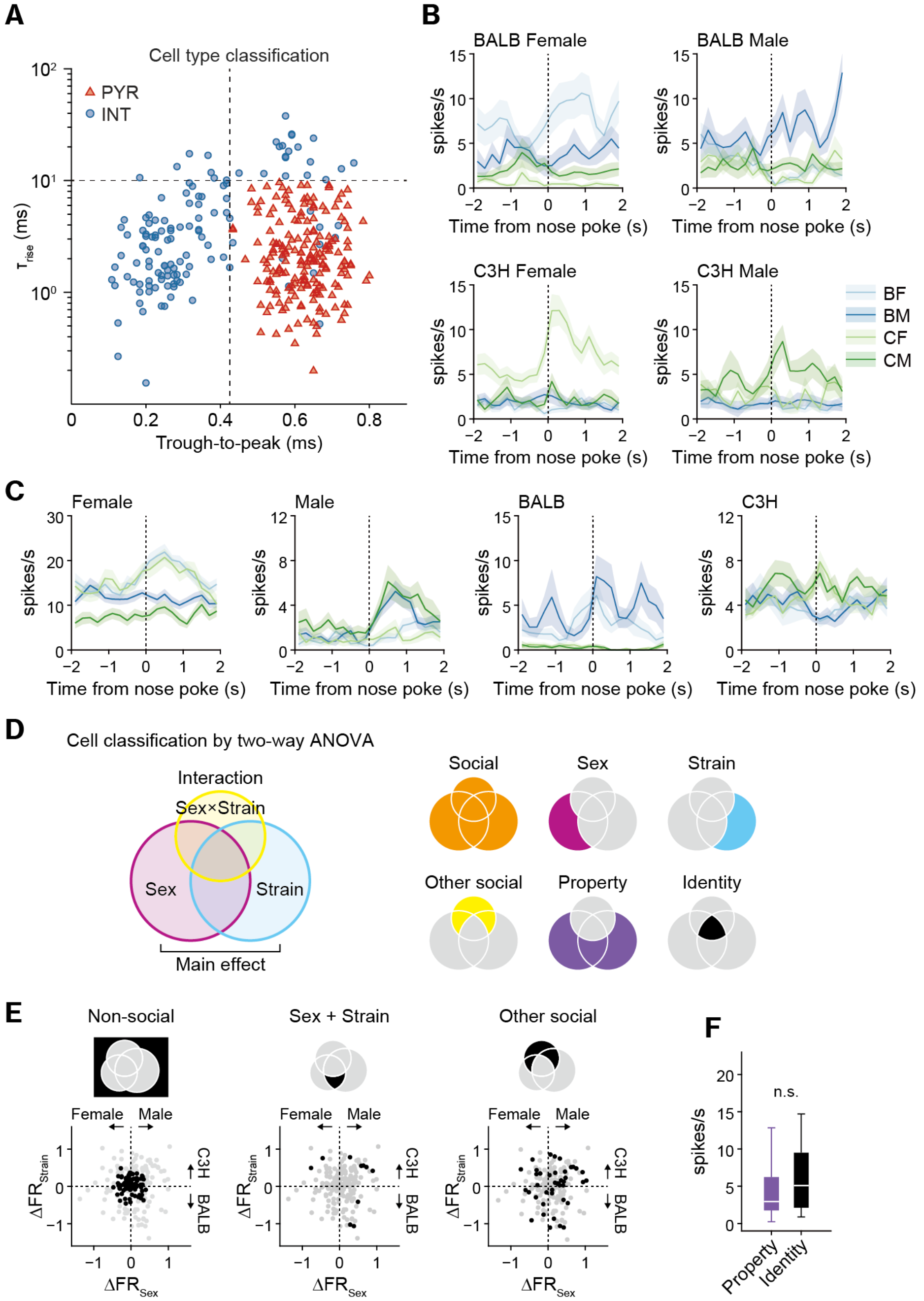


**fig. S4. Classification of vCA1 pyramidal cells.**

(**A**) Scatter plot of the classified units. Red triangles represent putative pyramidal cells (PYR) and blue circles represent interneurons (INT). (**B**, **C**) Peri-event time histogram of additional example social identity cells (B), sex and strain cells (C) aligned to the nose-poke onset. **(D)** Cells were classified using a two-way ANOVA, which analyzed the main effects of sex and strain of the social stimuli, as well as their interaction term. (**E**) Distribution of cell groups on a 2-D map showing differences in z-scored firing rates by sex and strain of social stimuli. (**F**) The firing rates of social property cells and identity cells. $P = 0.11$ by Wilcoxon rank-sum test.

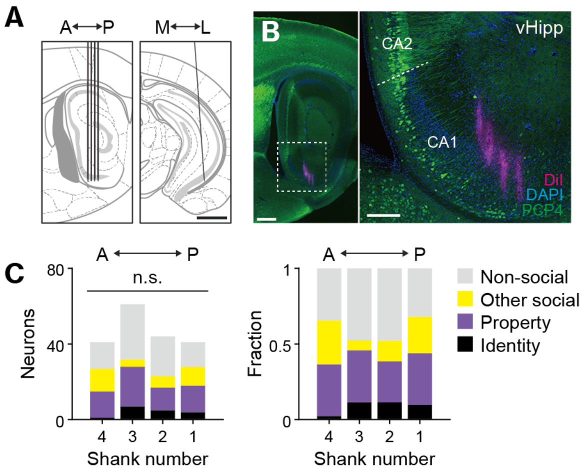


**fig. S5. Validation of electrode targeting vCA1.**

(**A**) Schematic of the probe implant location. Scale bar, 1 mm. (**B**) A representative image of the ventral hippocampus in the sagittal plane. The high-magnification image on the right is an enlargement of the white dashed square shown in the low-magnification image on the left, where the white dashed line marks the boundary between CA2 and CA1. Scale bar, 500 µm (left) and 200 µm (right). (**C**) The number (left) and a fraction (right) of recoded cells grouped by the probe shank along the anteroposterior axis. n.s., not significant; chi-square test of independence.

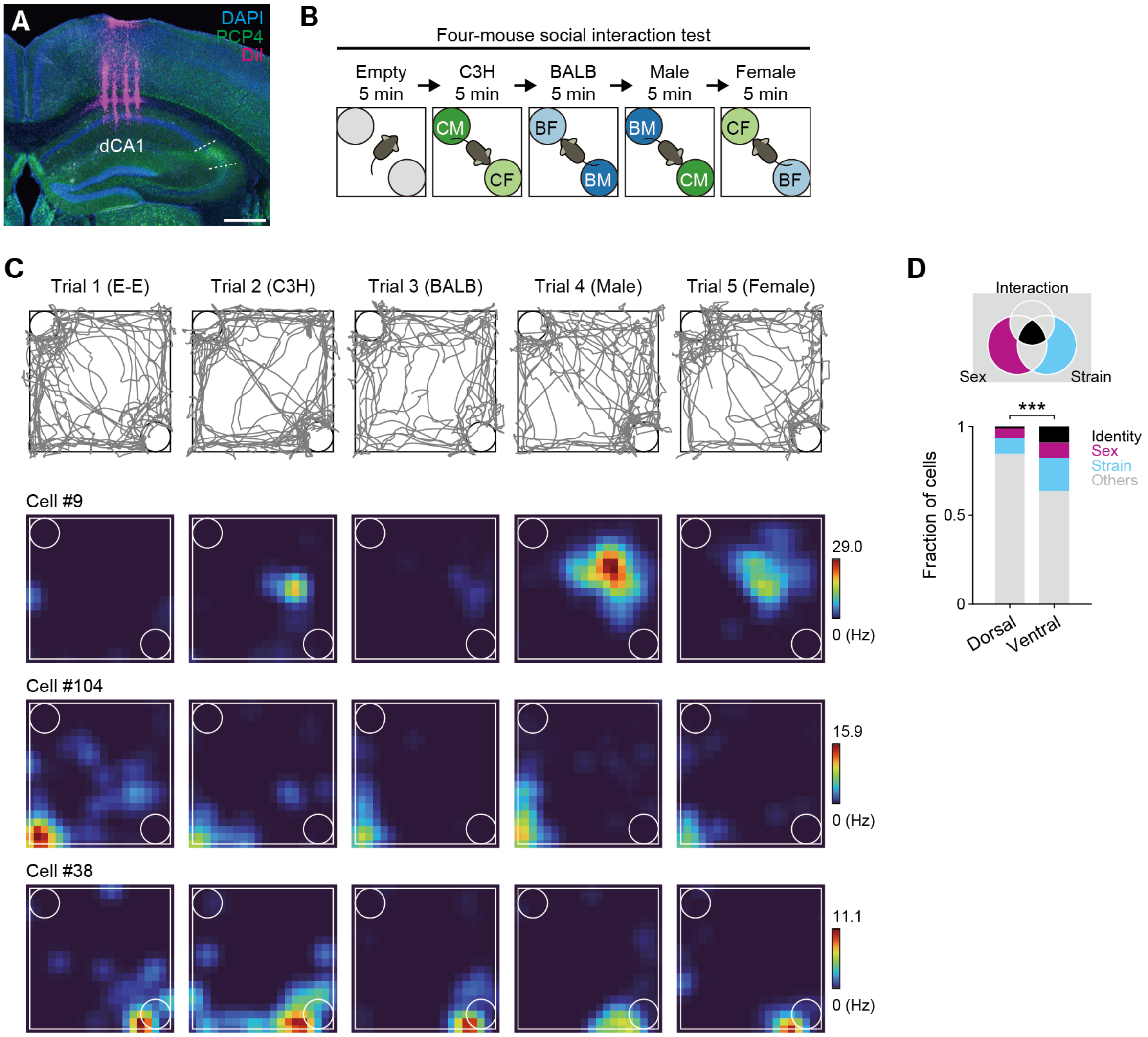


**fig. S6. Neural recordings from dCA1 during the four-mouse social interaction test.**

(**A**) Recording site with a scale bar of 500 µm. (**B**) Experimental paradigm for the four-mouse social interaction test. BF, BM, CF, and CM refer to BALB/c female, BALB/c male, C3H female, and C3H male mice, respectively. (**C**) Behavioral trajectories (top) and firing rate heatmaps of the three example cells. (**D**) Comparison of cell fractions between dCA1 and vCA1. ***$P < 0.001$, chi-square test of independence.

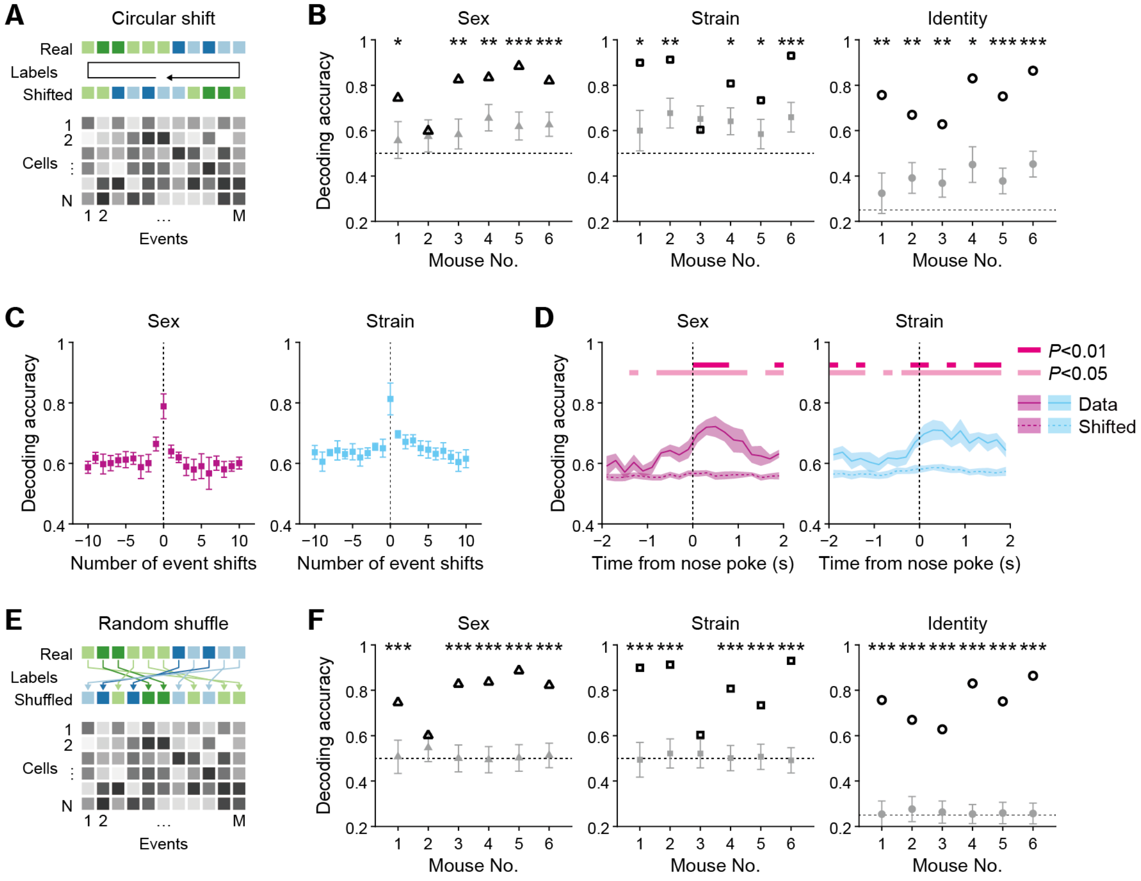


**fig. S7. Population decoding analysis.**

(**A**) Scheme for constructing null models using the circular shifting method. (**B**) Decoding accuracy of social identity, sex, and strain for each subject mouse. Error bars of the null models represent mean ± SD. The dotted lines represent chance labels. **P* < 0.05, ***P* < 0.01, ****P* < 0.001, determined by comparison to the null models. (**C**) Decoding accuracy of the classifier with real data (shifts = 0) and event-shifted labels (shifts = ±10 events) for identity. (**D**) Decoding accuracy for sex and strain over time (mean ± SEM, $n = 6$ mice). Solid lines with shaded areas represent the decoding accuracy computed from data with real labels, whereas dotted lines with shaded areas indicate the accuracy computed from data with circularly shifted labels. Significance was assessed by the Tukey–Kramer multiple comparisons. (**E**, **F**) Similar to (A) and (B), null models were constructed using the random shuffling method. ****P* < 0.001, determined by comparison to the null models.

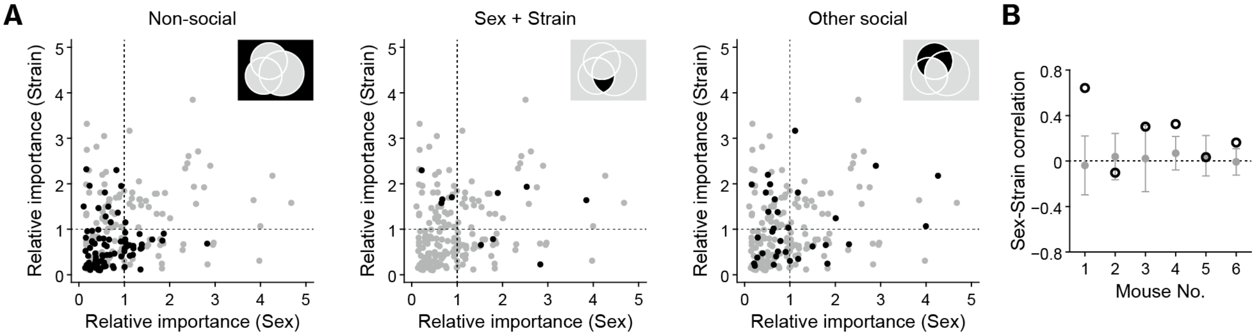


**fig. S8. Mixed selectivity analysis.**

(**A**) Scatter plot of the relative importance of sex decoding versus strain decoding for each cell group. (**B**) Correlation analysis repeated for each subject mouse. Error bars of the null models represent the mean ± SD.

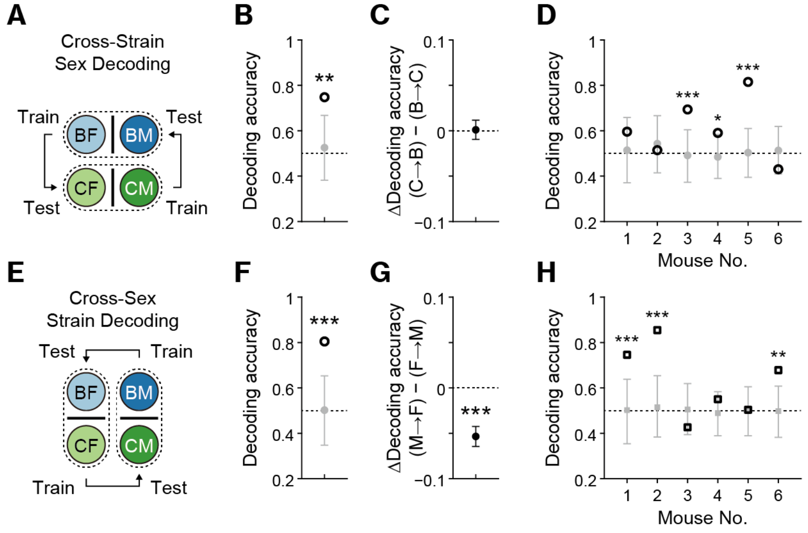


**fig. S9. Cross-condition generalization performance (CCGP).**

(**A**) Scheme for computing the sex CCGP. (**B**) Sex CCGP for the pseudo-population. $^{**}P < 0.01$, determined by comparison with the null model. The error bar for the null model represents the mean ± SD. (**C**) Comparison of the CCGPs trained on BALB/c data and tested on C3H data (B→C), and those trained on C3H data and tested on BALB/c data (C→B). (**D**) An analysis identical to (B) was conducted for each subject mouse. $^{*}P < 0.05$, $^{***}P < 0.001$, determined by comparison with the null models. (**E**) Scheme for computing the strain CCGP. (**F**) Strain CCGP for the pseudo-population. $^{***}P < 0.001$, determined by comparison with the null model. The error bar for the null model represents the mean ± SD. (**G**) Comparison of CCGPs trained on female data and tested on male data (F→M), and those trained on male data and tested on female data (M→F). $^{***}P < 0.001$, determined by the *t* test. (**H**) An analysis identical to (F) was conducted for each subject mouse. $^{**}P < 0.01$, $^{***}P < 0.001$, determined by comparison with the null models.

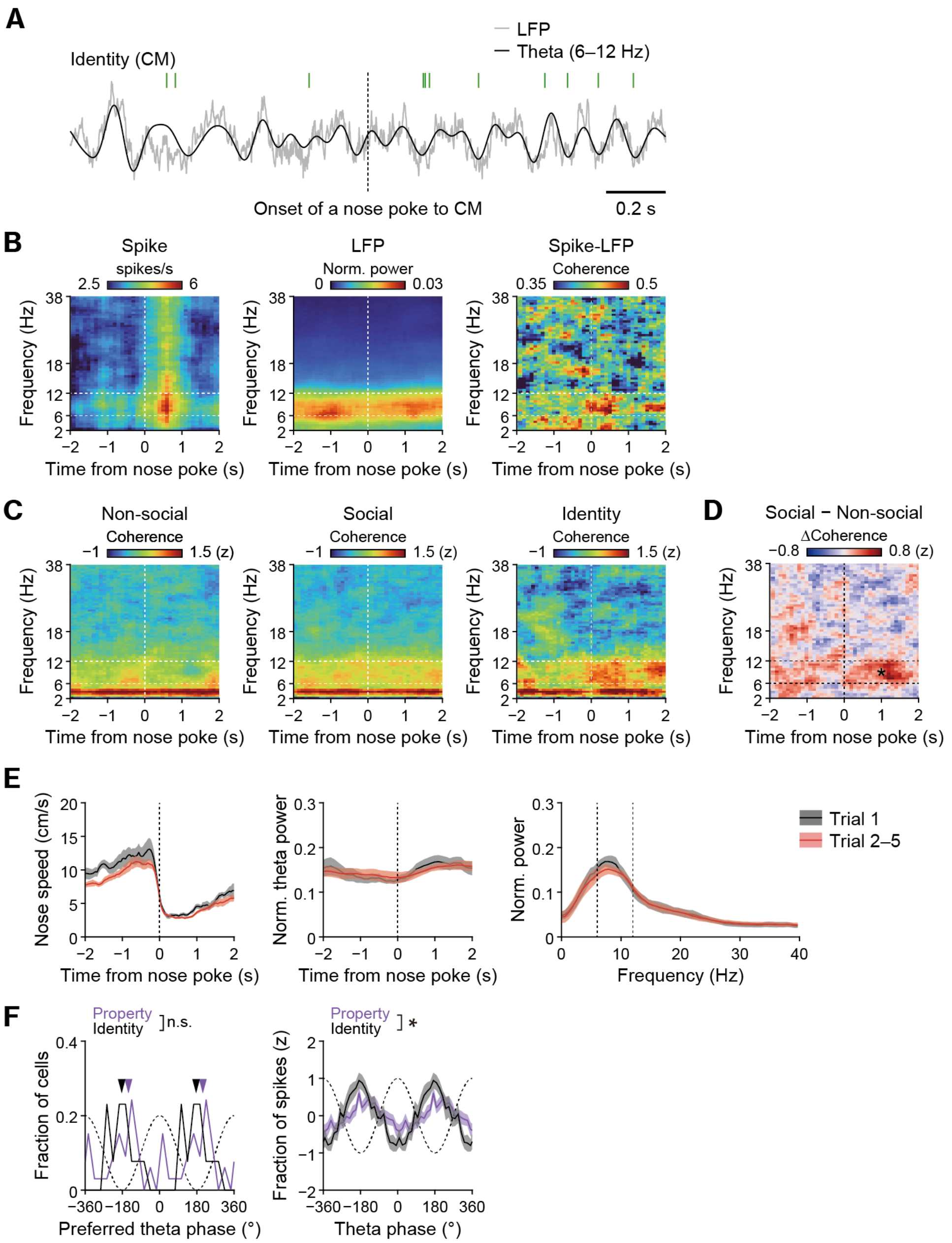


**fig. S10. Single cell theta modulation analysis.**

(**A**) LFP trace and spike timings for a representative neuron around the onset of social interaction, with spike timings marked by vertical green lines. (**B**) Spike spectrogram (left), LFP spectrogram (center), and spike-LFP coherogram (right) of the neuron in (A) around the onset of social interaction. Note the absence of an increase in instantaneous theta power during the interaction. (**C**) Z-scored spike-LFP coherograms averaged within non-social, social, and social identity cells. (**D**) Comparison of average spike-LFP coherograms between social cells and non-social cells. **P* < 0.05, permutation test. (**E**) Left and center, speed of the subject mice's nose and normalized theta power aligned with nose-poke onset. Right, normalized LFP power during social interactions.

Vertical dotted lines indicate the theta frequency range (6–12 Hz). (**F**) Left, distribution of preferred theta phases for social property cells and social identity cells. The arrowheads indicate the average preferred phase for each group. n.s., not significant, Watson–Williams test. Right, mean z-scored proportions of spike theta phases for social property cells and social identity cells. *$P < 0.05$, permutation test.

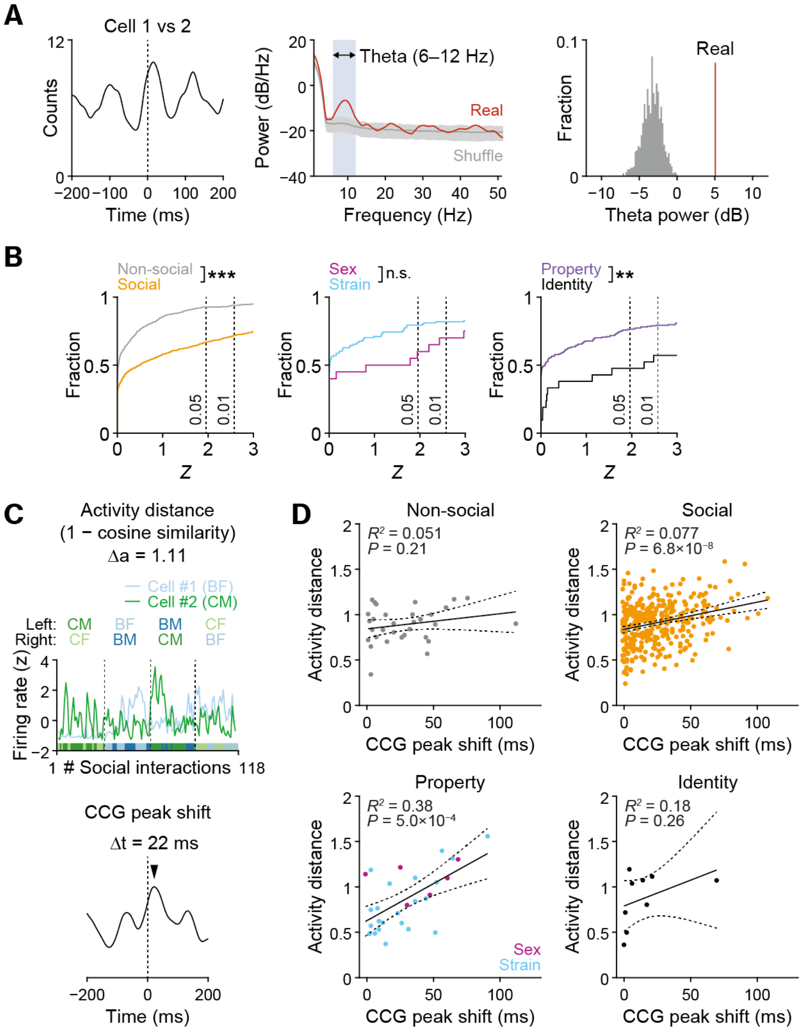


**fig. S11. Cell-pair theta modulation analysis.**

(**A**) Method for determining the significance of the CCG theta modulation. (**B**) The cumulative density function of theta modulation z-scores derived from the *P* values of the permutation test described above. Vertical dotted lines indicate the corresponding *P* values. $**P < 0.01$ and $***P < 0.001$, Kolmogorov–Smirnov test. (**C**) Top, response series to social stimuli across trials for two examples simultaneously recorded social identity cells, with cell #1 preferring BF (light blue line) and cell #2 preferring CM (dark green line). The color code at the bottom indicates the social stimulus target in each interaction, where light and dark blue represent BF and BM, respectively, and light and dark green represent the CF and CM, respectively. The vertical dotted lines indicate the borders between the trials. Bottom, spike cross correlogram (CCG) for the same cell pair, shown in arbitrary units. (**D**) Correlation between the temporal shift in CCG peaks and the pairwise distance in neuronal activity during social interactions for pairs of non-social ($n$ = 33), social ($n$ = 365), sex and strain ($n$ = 28), and identity ($n$ = 9) cell pairs.

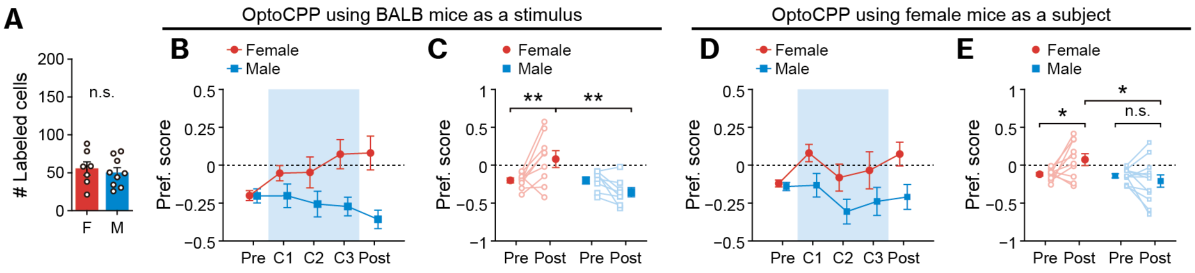


**fig. S12. OptoCPP test under additional conditions.**

(**A**) Number of EYFP-expressing cells labeled by social interactions with a female (F; $n = 7$ mice) or a male (M; $n = 9$ mice) stimulus mouse. (**B**) Temporal dynamics of preference scores over conditioning. $n$= 9 (female) and 8 (male) mice. (**C**) Comparison of preference scores between the Pre-test and Post-test. $**P < 0.01$, Tukey–Kramer multiple comparisons. (**D**) Temporal development of preference scores during conditioning. $n = 10$ (Female) and 12 (Male) mice. (**E**) Comparison of the preference scores between the Pre-test and Post-test. $*P < 0.05$, Tukey–Kramer multiple comparisons.

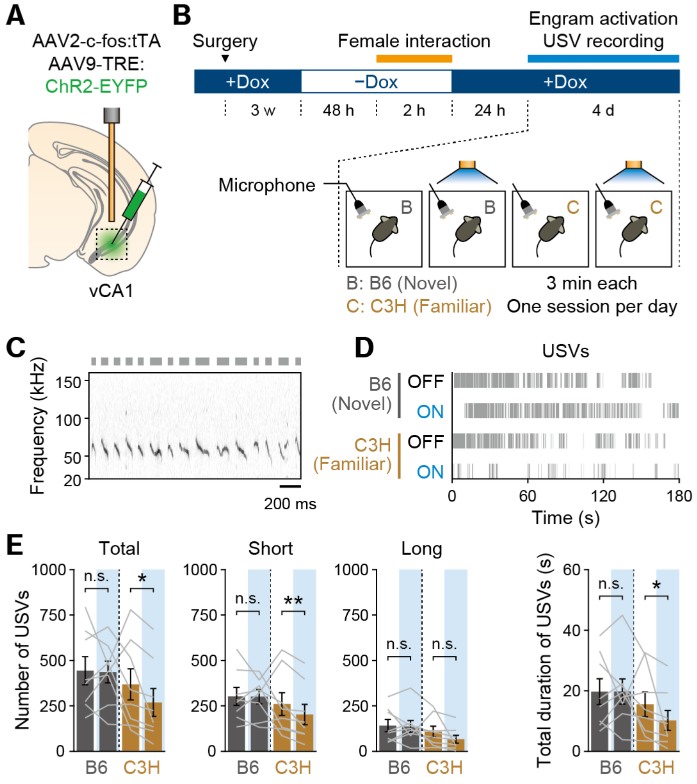


**fig. S13. USV recordings during female engram activation.**

(**A**) Schematic representation of virus-mediated female engram labeling. (**B**) Experimental paradigm for female engram labeling and activation along with USV recording in the presence of a female mouse ($n$ = 6 mice). (**C**) Representative spectrogram during the interaction with a novel B6 female mouse. The thick gray line above the spectrogram indicates the periods of detected USV syllables. (**D**) Representative raster plots showing USV syllables when the subjects were presented with a B6 (novel) or a C3H (familiar) female mouse, with or without optogenetic activation of the female engram. (**E**) Number and total duration of USVs, with the number classified as short (3–50 ms) or long (50–300 ms) syllables, and the total duration calculated as their sum. *$P$ < 0.05, **$P$ < 0.01, Tukey–Kramer multiple comparisons.

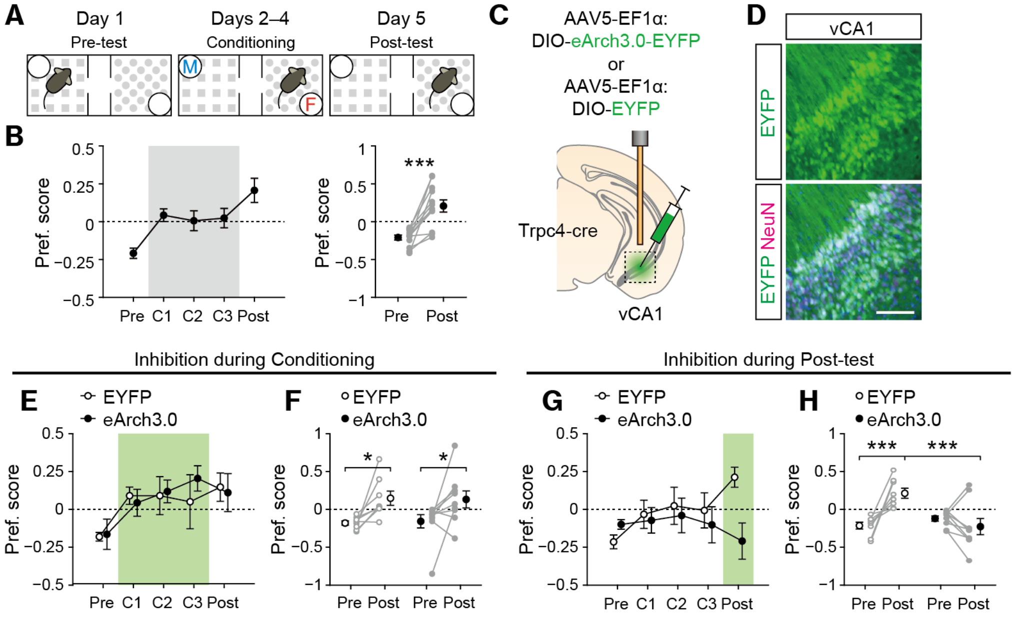


**fig. S14. The distinct roles of vCA1 in real-time and conditioned place preference induced by live female mice.**

(**A**) Experimental protocol for conditioned place preference test using live mice. (**B**) Left: temporal dynamics of preference scores during conditioning ($n$ = 12 mice). Right: comparison of the preference scores between the Pre-test and Post-test. ***$P < 0.001$, paired $t$ test. (**C**) Schematic representation of optogenetic inhibition targeting vCA1. (**D**) Representative images of viral expression. Scale bar, 100 µm. (**E**, **F**) Temporal dynamics (E) and comparison between Pre-test and Post-test (F) of preference scores with vCA1 optogenetic inhibition during conditioning. $n$ = 8 (EYFP) and $n$ = 9 (eArch3.0) mice. *$P < 0.05$, Tukey–Kramer multiple comparisons. (**G**, **H**) Similar to (E) and (F), vCA1 was inhibited during the Post-test. $n$ = 8 (EYFP) and $n$ = 10 (eArch3.0) mice. ***$P < 0.001$, Tukey–Kramer multiple comparisons.

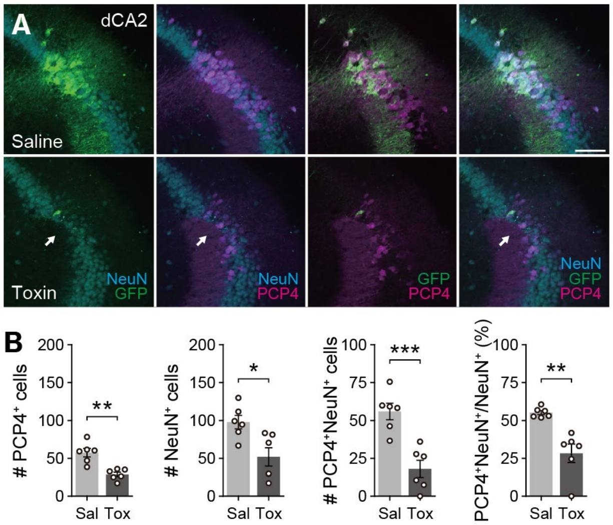


**fig. S15. Quantification of dCA2 ablation.**

(**A**) Representative images of dCA2. The images shown here are the same as those presented in Fig. 4F but displayed using different color combinations to highlight specific staining. Scale bar, 100 µm. (**B**) Number of $PCP4^+$ cells, number of $NeuN^+$ cells, number of $PCP4^+NeuN^+$ double-positive cells, and ratio of double-positive cells to $NeuN^+$ cells. *$P < 0.05$, **$P < 0.01$, ***$P < 0.001$, Student's *t* test.

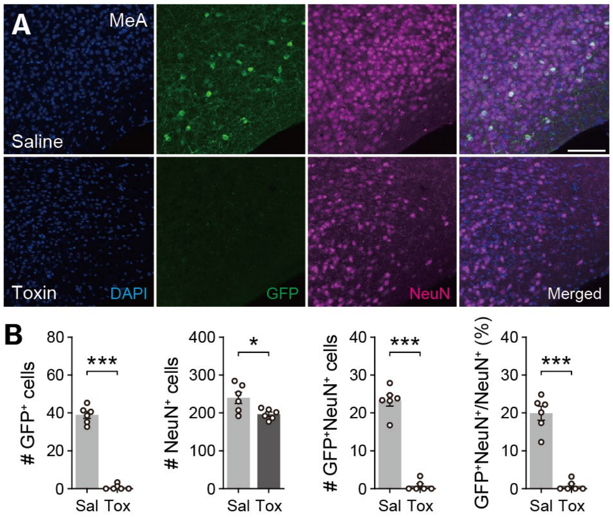


**fig. S16. Quantification of MeA ablation.**

(**A**) Representative images of the MeA. Scale bar, 100 μm. (**B**) Number of $GFP^+$ cells, number of $NeuN^+$ cells, number of $GFP^+NeuN^+$ double-positive cells, and ratio of double-positive cells to $NeuN^+$ cells. *$P < 0.05$, ***$P < 0.001$, Student's *t* test.

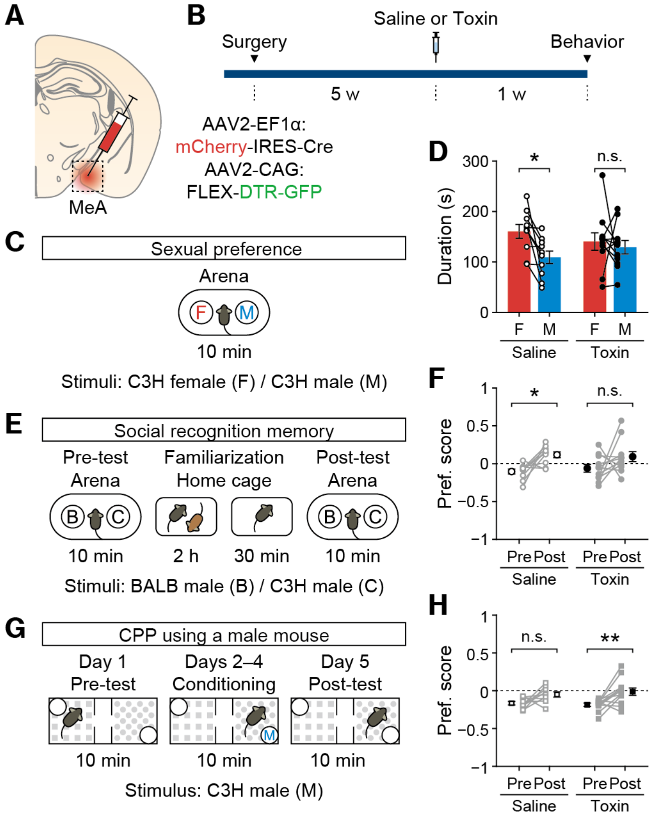


**fig. S17. Behavioral tests after MeA ablation.**

(**A**) Schematic representation of MeA ablation using a Cre-inducible DTR. (**B**) Timeline of behavioral tests following MeA ablation. (**C**) Experimental design of the sexual preference test. (**D**) Total duration of dwelling time spent in the social zone around female or male social stimuli during the sexual preference test. *$P < 0.05$, determined by Tukey–Kramer multiple comparisons. (**E**) Experimental design of the social recognition test. (**F**) Preference score between BALB/c and C3H male mouse during the Pre-test and Post-test of the social recognition memory test. *$P < 0.05$, Tukey–Kramer multiple comparisons. (**G**) Experimental design of CPP using a male mouse. (**H**) Preference scores for the conditioned side during the Pre-test and Post-test. **$P < 0.01$, Tukey–Kramer multiple comparisons.

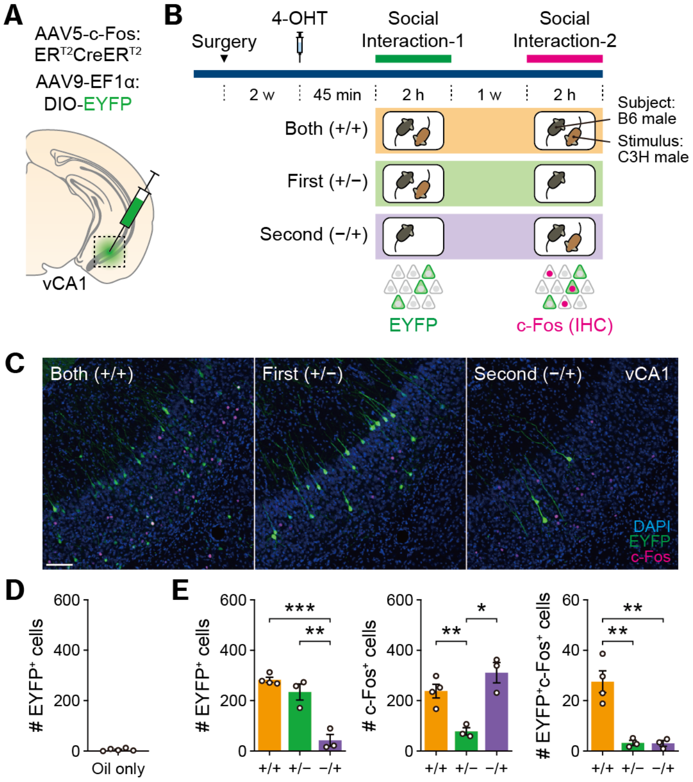


**fig. S18. Validation of the c-fos:ER$^{T2}$CreER$^{T2}$ construct.**

(**A**) Schematic representation of virus cocktail injection. (**B**) Experimental paradigm fior activity-dependent cell labeling. (**C**) Representative images of vCA1 in each group. Scale bar, 100 µm. (**D**) The number of EYFP$^+$ cells following social interactions in the absence of 4-OHT. It should be noted that virtually no cell expressed EYFP. (**E**) Number of EYFP$^+$ cells, number of c-Fos$^+$ cells, and number of EYFP$^+$c-Fos$^+$ double-positive cells. *$P < 0.05$, **$P < 0.01$, ***$P < 0.001$, Tukey–Kramer multiple comparisons.

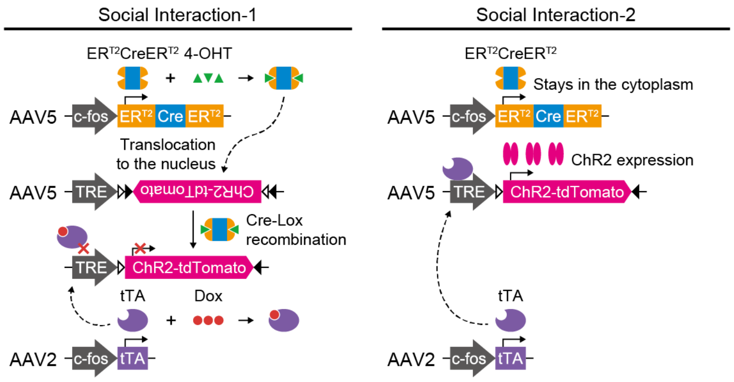


**fig. S19. Intersectional labeling of dual-activated neurons.**

Mice received bilateral vCA1 injections of AAV9-TRE:DIO-ChR2-tdTomato, AAV5-c-fos:ER$^{T2}$CreER$^{T2}$-PEST, and AAV2-c-fos:tTA. During the first social interaction, ER$^{T2}$CreER$^{T2}$ is expressed in a *c-fos*-dependent manner, and the presence of 4-hydroxytamoxifen (4-OHT) enables its translocation to the nucleus. This process triggers the Cre-Lox recombination of the ChR2-tdTomato sequence. Subsequently, the second social interaction, conducted in the absence of doxycycline (Dox), induces *c-fos*-dependent activation of the tTA/TRE system. This resulted in ChR2 expression exclusively in neurons that were activated during both social interactions.

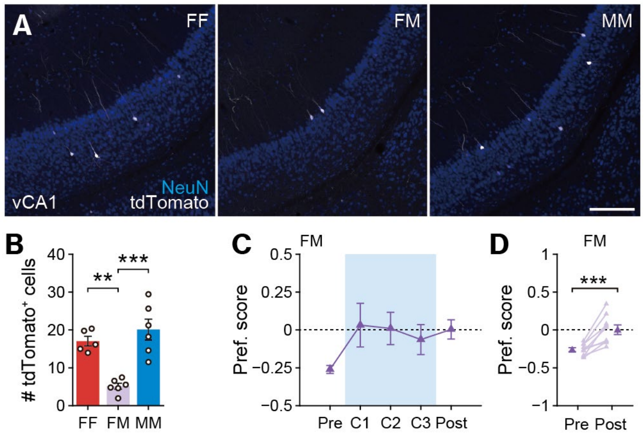


**fig. S20. Intersectional labeling and manipulation of neurons activated by different sex.**

(**A**) Representative images of vCA1 following intersectional cell labeling. Scale bar, 200 µm. (**B**) The number of tdTomato$^{+}$ cells in each group. ***P* < 0.01, ****P* < 0.001, Tukey–Kramer multiple comparisons. (**C**) Temporal dynamics of preference scores during conditioning. (**D**) Comparison of preference scores between the Pre-test and Post-test. ****P* < 0.001, paired *t* test.
